# Engineering Programmable Material-To-Cell Pathways Via Synthetic Notch Receptors To Spatially Control Cellular Phenotypes In Multi-Cellular Constructs

**DOI:** 10.1101/2023.05.19.541497

**Authors:** Mher Garibyan, Tyler Hoffman, Thijs Makaske, Stephanie Do, Alexander R March, Nathan Cho, Nico Pedroncelli, Ricardo Espinosa Lima, Jennifer Soto, Brooke Jackson, Ali Khademhosseini, Song Li, Megan McCain, Leonardo Morsut

**Affiliations:** Department of Stem Cell Biology and Regenerative Medicine, Keck School of Medicine of USC, University of Southern California; Eli and Edythe Broad Center; Alfred E. Mann Department of Biomedical Engineering, USC Viterbi School of Engineering, University of Southern California; Department of Bioengineering, University of California Los Angeles, Los Angeles, CA, USA

## Abstract

Synthetic Notch (synNotch) receptors are modular synthetic components that are genetically engineered into mammalian cells to detect signals presented by neighboring cells and respond by activating prescribed transcriptional programs. To date, synNotch has been used to program therapeutic cells and pattern morphogenesis in multicellular systems. However, cell-presented ligands have limited versatility for applications that require spatial precision, such as tissue engineering. To address this, we developed a suite of materials to activate synNotch receptors and serve as generalizable platforms for generating user-defined material-to-cell signaling pathways. First, we demonstrate that synNotch ligands, such as GFP, can be conjugated to cell-generated ECM proteins via genetic engineering of fibronectin produced by fibroblasts. We then used enzymatic or click chemistry to covalently link synNotch ligands to gelatin polymers to activate synNotch receptors in cells grown on or within a hydrogel. To achieve microscale control over synNotch activation in cell monolayers, we microcontact printed synNotch ligands onto a surface. We also patterned tissues comprising cells with up to three distinct phenotypes by engineering cells with two distinct synthetic pathways and culturing them on surfaces microfluidically patterned with two synNotch ligands.

We showcase this technology by co-transdifferentiating fibroblasts into skeletal muscle or endothelial cell precursors in user-defined spatial patterns towards the engineering of muscle tissue with prescribed vascular networks. Collectively, this suite of approaches extends the synNotch toolkit and provides novel avenues for spatially controlling cellular phenotypes in mammalian multicellular systems, with many broad applications in developmental biology, synthetic morphogenesis, human tissue modeling, and regenerative medicine.

## Main

A fundamental goal for the emerging area of synthetic morphogenesis and tissue engineering is the ability to design and spatially control gene expression patterns within a multicellular construct. Intricate patterns of gene expression control the proper organization and physiology of cells, tissues, and organs and are a hallmark of complex multicellular systems across the tree of life. Individual cells express genetic networks that drive or support cell fate commitment and functional behaviors, like motility and proliferation. During embryonic development, initially uniform cell ensembles activate genetic networks in designated spatial regions to generate tissues with distinct geometrical patterns. The spatial organization of cells within a tissue endows them to coordinate and accomplish complex functions, such as absorption or contractility. *In vivo*, spatial domains of gene expression are driven by genetically-encoded communication networks involving intra-cellular^1, 2^, inter-cellular^3–7^, and cell-to-extracellular matrix (ECM) and ECM-to-cell^8–10^ components. Several of these networks are active in organoids *in vitro*, which self-organize and replicate select microscale architectural features similar to native tissues. However, genetic networks in organoids are spatially activated in an autonomous way and some genetic networks fail to activate at all, leading to heterogeneity and stunted tissue structures. Because self-organization is convoluted with differentiation and other complex cell behaviors, *in vitro* methods for arbitrarily engineering and interrogating spatial gene expression patterns and their impact would augment our understanding of biological systems^11–14^. Advanced technologies for spatially controlling gene expression would also enable tissues to be engineered with user-defined cellular compositions and geometries, which would be impactful for the fields of regenerative medicine, Organs on Chips, and lab-grown protein-rich food sources^15–18^.

Classically, tissue engineers have focused on influencing cell differentiation and behavior by engaging endogenous cell surface receptors. For example, natural ligands, such as extracellular matrix proteins, can be presented to cells in user-defined spatial arrangements via microfabricated biomaterials to control adhesion, alignment, or differentiation^19–21^. Because these approaches rely on engagement of endogenous receptors, such as integrins, stereotyped and often complex behaviors are activated in responding cells. However, with these approaches, users are confined to the limited library of endogenous ligands and receptors and their pre-existing downstream pathways, many of which are multifaceted with ambiguous outcomes. Recently, synthetic receptors have been developed that endow cells with orthogonal, customizable signaling capabilities^22^. Thus, we reasoned that these receptors could be leveraged to spatially control gene expression patterns in engineered tissues with more precision than endogenous receptors. Specifically, we turned to a class of synthetic receptors based on native Notch signaling, named synthetic Notch or synNotch^23^. SynNotch are a class of synthetic receptors composed of chimeric protein domains: an antibody-based binding extracellular domain (e.g. anti-GFP nanobody), the Notch juxtamembrane and transmembrane domains, and orthogonal transcription factors (e.g. Gal4) as the intracellular domain. SynNotch receptors have many desirable features that could be exploited to spatially control gene expression: (i) the receptor is not activated by soluble factors; (ii) the ligand is customizable and can be an orthogonal inert molecule, such as a fluorescent protein (e.g. GFP); (iii) receptor activation can drive customizable cellular responses, such as differentiation, when combined with complementary genetically engineered cassettes.

SynNotch has previously been used to generate spatial patterns of gene expression in 2D (concentric rings^23^) and in 3D (polarized and layered spheroids^24^) by using neighboring cells (i.e., sender cells) to present synthetic ligands to cells expressing synNotch (i.e., receiver cells). Cellular ligand presentation, however, has the disadvantage that controlling the geometry of synthetic ligands necessitates controlling the location of sender cells, making the problem circular. Evidence suggests that a pulling force between sender and receiver cells is necessary to initiate signal transduction in the receiving cell, similar to endogenous Notch receptors. Due to this feature, synNotch has also been activated by synthetic ligands passively adsorbed onto cell culture surfaces^23^, tethered by DNA linkers to microbeads^25^, attached to atomic force microscopy probes^26^, or conjugated to the surfaces of biomaterials^27^. However, none of these approaches attempted patterning of ligands, a feature that would enable spatial control of synNotch activation and therefore gene expression.

Here, our objective was to develop generalizable, user-defined, material-to-cell pathways for spatially controlling genetic networks and differentiation in multicellular constructs via synNotch. We first show that we can activate synNotch with synthetic ligands (e.g., GFP) presented by materials that offer increasing degrees of spatial control: (i) genetically encoded, cell-produced extracellular matrix proteins (e.g., fibronectin-GFP fusions), (ii) extracellular matrix-derived hydrogels, and (iii) microcontact printed culture surfaces. We also show that these approaches are generalizable to multiple synNotch receptors and can activate distinct synthetic pathways in cells with two synNotch receptors (i.e., dual-receiver cells). We then show that these approaches can be extended to spatially control patterns of gene expression and cell fate by transdifferentiating embryonic fibroblasts into either skeletal muscle precursors or endothelial cell precursors in tissue-relevant geometries. Finally, we demonstrate a method for spatially controlling the co-transdifferentiation of fibroblasts to one of two cell fates (endothelial cell precursors or skeletal muscle precursors) in a continuous tissue construct. This was achieved by generating dual-lineage fibroblasts expressing two independent synNotch receptors (one for endothelial transdifferentiation, one for muscle transdifferentiation) and culturing these cells on a surface with the two synthetic cognate ligands patterned via a microfluidic device. These novel methods for generating spatial patterns of gene expression and cell fate add a powerful and flexible functionality to the synthetic biology toolbox for controlling and investigating multicellular organization.

### Activation of synNotch from particles and cell-generated ECM

To evaluate the activation of synNotch receptors by synthetic ligands presented on materials, we first used a suspension of microparticles to present ligands semi-analogously to the presentation of ligands on the membranes of sender cells. We tethered GFP to carboxyl-modified microparticles of different diameters (2um-10um) using an EDC/NHS reaction. This approach enables different amounts of GFP to be loaded by simply adjusting the concentration of GFP in the conjugation reaction. We then added these microparticles to a monolayer of receiver fibroblasts (L929 cells) that were engineered to express an anti-GFP/tTA synNotch receptor and its mCherry reporter gene (Figure 1A). As expected, mCherry fluorescence at 24h post-seeding increased with the concentration of GFP loaded onto the microparticles for all particle diameters and was absent when cells were presented with unmodified particles (Figure 1B-C and Fig. S1A-B). Importantly, 5um microparticles loaded with 500 or 1000 ug/mL GFP induced mCherry in the receiver fibroblasts at a level similar to GFP-presenting sender cells co-cultured at a 1:1 ratio, indicating that synthetic ligands conjugated to microparticles can activate synNotch receptors to a similar extent as synthetic ligands presented by sender cells.

**Figure 1.**
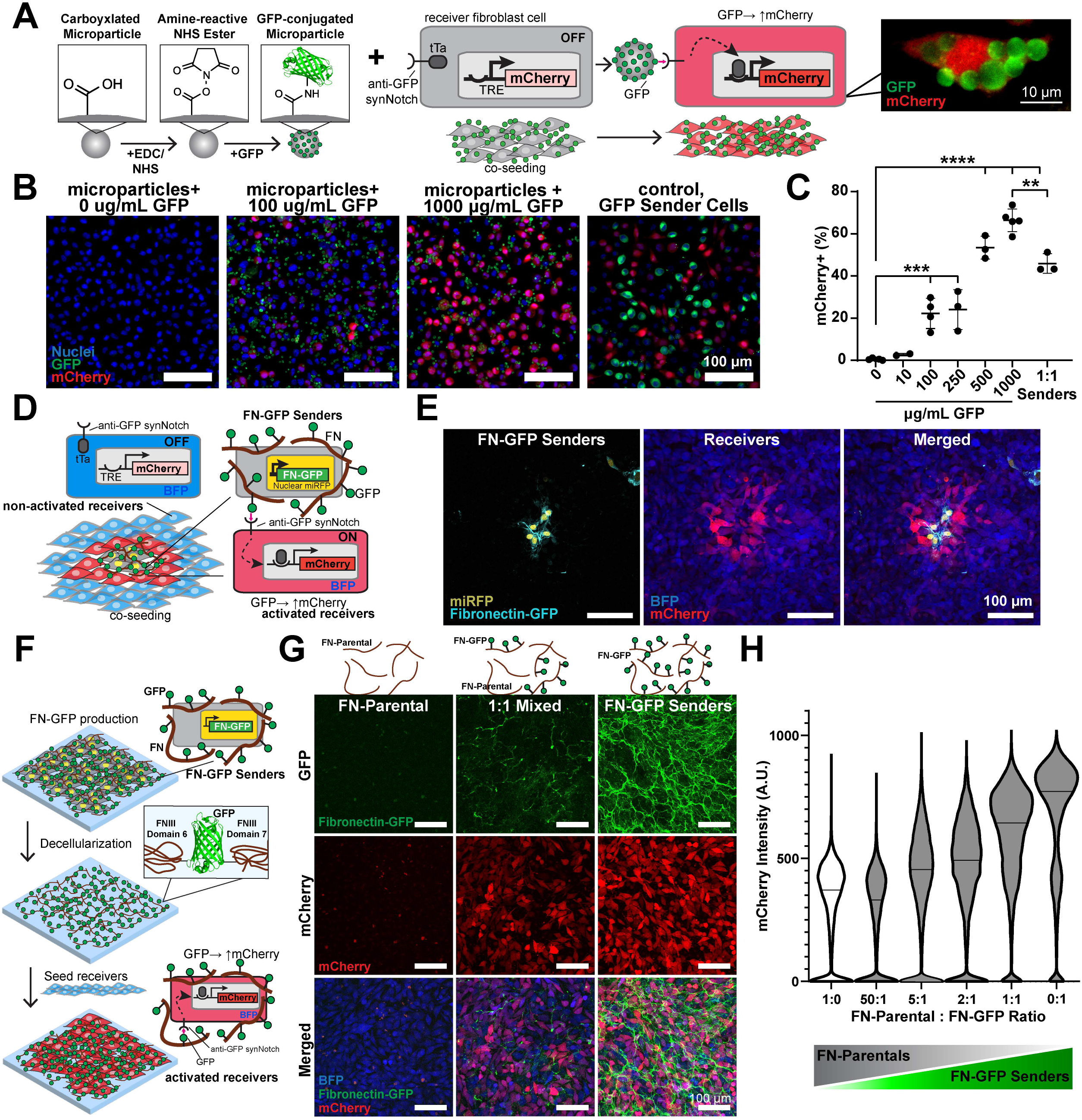
Microparticle-conjugated GFP and cell-deposited fibronectin-GFP activate synNotch. (A) Schematic of GFP conjugation to microparticles and subsequent co-culture with anti-GFP synNotch→mCherry fibroblasts; fluorescence image of an individual activated receiver fibroblast in the presence of GFP-loaded-microparticles. (B) Fluorescence microscopy images of synNotch receiver fibroblasts cultured in the presence of 0, 100, and 1000 ug/mL 5pm GFP- conjugated microparticles or GFP-presenting sender cells following 24 hours culture. Scale bars, 100pm. (C) Percent of mCherry expressing cells quantified by image analysis following 24-hour co-culture with GFP microparticles or GFP sender cells. Data represents mean±s.d, n=3-5, p<0.01(**), p<0.001(***), p<0.0001(****). (D) Schematic of Fibronectin-GFP (FN-GFP) producing fibroblasts, with an miRFP nuclear tag co-cultured with anti-GFP synNotch^mCherry fibroblast that constitutively express a BFP reporter. (E) Fluorescence microscopy images of co-cultured FN-GFP cells and synNotch receiver fibroblasts taken after 72 hours of culture. Scale bars, 100pm. (F) Schematic of FN-GFP deposition by NF-GFP senders, decellularization, and subsequent reseeding with synNotch fibroblasts. (G) Top row: schematics of decellularized extracellular matrix (ECM) present on the surface plates before seeding, which is produced by 8 days of culture of different ratios of parental fibroblasts (FN-Parental) and FN-GFP senders (1:0, 1:1 or 0:1, FN-Parental:FN-GFP) prior to decellularization. Bottom: microscope images of anti- GFP synNotch fibroblasts 48 hours following seeding onto the corresponding decellularized ECM. Scale bars, 100pm. (H) Flow cytometry mCherry expression of synNotch fibroblasts seeded onto decellularized ECM prepared from varying ratios of FN:Parental and FN-GFP fibroblasts, taken 48 hours following synNotch fibroblast seeding. Arbitrary Units (A.U.). Data represents distribution of individual cell intensity and median value.

We next asked if synNotch receptors could be activated by synthetic ligands presented on extracellular matrix fibers produced by cells. Thus, we genetically engineered mouse embryonic fibroblasts (3T3 cells) to produce a fusion protein of fibronectin and GFP (FN-GFP^28^). These cells were also engineered to express a far-red fluorescent nuclear reporter protein. We hypothesized that these FN-GFP sender cells would deposit an extracellular matrix containing synthetic ligands that would signal to receiver cells. To test this, we co-cultured a low amount of FN-GFP sender cells alongside receiver fibroblasts expressing anti-GFP/tTA synNotch receptors that activate mCherry (Fig. 1D). At 72h after seeding, we observed mCherry expression only in receiver cells that are near to FN-GFP sender cells (Fig. 1E), indicating that the anti-GFP synNotch receptor is locally activated in response to FN-GFP embedded in the extracellular matrix.

We then tested if cell-deposited FN-GFP matrices can activate receiver cells after the sender cells are removed. To do so, we cultured FN-GFP sender cells as a monolayer for 8 days and subsequently performed decellularization to remove all cellular components while preserving the extracellular matrix (see Fig. 1F and S1C-D). Receiver cells cultured on the decellularized matrices for 48h expressed mCherry, indicating that synthetic ligands embedded in the extracellular matrix remained functional through the decellularization process. To tune the level of synNotch receptor activation by decellularized matrices, we co-cultured FN-GFP sender cells with the unmodified parental 3T3 cells at various ratios. We similarly decellularized the co-cultured tissues and then seeded the decellularized matrices with receiver cells. mCherry intensity scaled with the ratio of parental cells to FN-GFP sender cells in the original tissue (Figure 1G-H), demonstrating tunability of activation of synNotch via cell-produced extracellular matrix fibers.

One advantage of synthetic receptors is that they can be engineered to both recognize distinct input ligands and drive distinct cellular responses. This feature has been used to generate a library of orthogonal synNotch receptors and pathways that function independently from each other and from endogenous receptors and pathways^23, 24, 29, 30^. For example, synNotch receptors have been developed to recognize mCherry as its ligand in^31^. To test if activation of synNotch receptors by matrix-presented synthetic ligands is generalizable to other ligand-receptor pairs, we generated FN-mCherry sender cells as well as corresponding receiver cells with anti-mCherry synNotch/Gal4 receptors that induce a BFP reporter gene upon activation. Similar to FN-GFP sender cells, FN-mCherry sender cells activate receiver cells in co-culture and upon decellularization (Fig. S1E-H). We also observed that anti-mCherry receiver cells were not activated by FN-GFP decellularized matrices, illustrating the orthogonality of receptor activation by matrix-presented synthetic ligands (Fig. S1I-J). Overall, these data demonstrate that synNotch receptors can be robustly, tunably, and modularly activated by ligands presented on cell-produced extracellular matrix fibers.

### Activation of synNotch from hydrogels in 2D and 3D

To improve user control and tunability, we next tested if synNotch could be activated by ligands presented on purified extracellular matrix fibers processed into hydrogel biomaterials. As a first step, we attempted to activate synNotch receptors in cells cultured on the surface of matrix-derived hydrogels. Based on our previous protocols^32, 33^, we fabricated slabs of gelatin hydrogels enzymatically cross-linked with transglutaminase, an enzyme that cross-links glutamine and lysine residues^34^. We next sought to conjugate GFP onto the hydrogel surface with transglutaminase by adapting methods for conjugating laminin onto gelatin^35^. However, GFP is weakly susceptible to transglutaminase because the glutamine and lysine residues of globular proteins are relatively inaccessible^36, 37^. Thus, we synthesized GFP with a short C terminus LACE peptide tag (GFP-LACE) to provide accessible lysine residues^38^. We then treated gelatin hydrogels with a solution of GFP-LACE and transglutaminase to conjugate GFP onto the surface (Fig. 2A)^35^. When receiver cells with anti-GFP synNotch/tTA receptors that activate miRFP were cultured on the GFP-gelatin hydrogels, miRFP intensity increased in a GFP dose-dependent manner (Fig. 2B-C). Thus, synNotch receptors can be activated by synthetic ligands presented on the surface of matrix-derived hydrogels.

**Figure 2.**
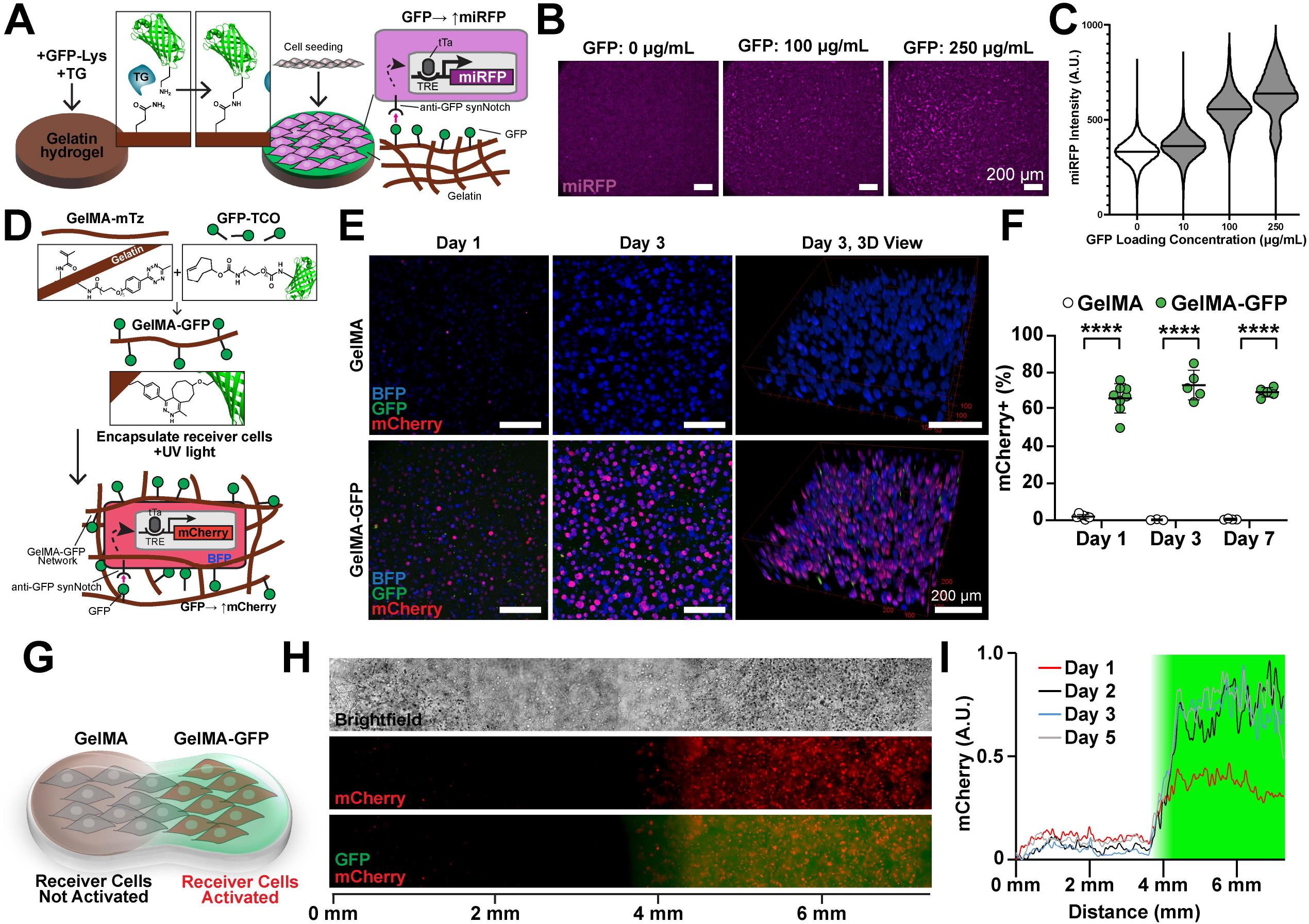
Hydrogel-conjugated ligands robustly and spatially activate reporter transgenes via synNotch. (A) Schematic of enzymatic GFP conjugation to surface of gelatin hydrogel using transglutaminase (TG) and resulting interactions with synNotch receiver fibroblasts. (B) Fluorescence microscopy images of anti-GFP/tTA synNotch receiver fibroblasts seeded on gelatin hydrogels conjugated with GFP at 0, 100 or 250 µg/mL. Scale bars, 200μm. (C) Flow cytometry analysis of miRFP expression of synNotch fibroblasts 72 hours following seeding onto GFP-conjugated hydrogel surfaces prepared with varying concentrations of GFP (0, 10, 100, 250 µg/mL). Data represents distribution of individual cell intensity and median value. (D) Schematic of gelatin methacryloyl-methyltetrazine (GelMA-mTz) covalent conjugation to GFP-trans-Cyclooctene (GFP-TCO) and subsequent engineered cell photoencapsulation with UV light. (E) Z-projected and 3D view of fluorescence microscopy images of anti-GFP synNotch receiver fibroblasts encapsulated within GelMA-mTz hydrogels containing 0 (GelMA) or 50μg/mL GFP- TCO (GelMA-GFP) at Day 1 and 3. Scale bars, 200μm. (F) Percent of mCherry expressing cells quantified by image analysis following 1, 3, and 7 days of culture within GelMA or GelMA-GFP hydrogels. Data represents mean ± s.d, n=3-8, p<0.0001(****). (G) Schematic of engineered cell encapsulation within bi-phasic hydrogel containing spatially localized GFP (H) Brightfield and fluorescence images of anti-GFP synNotch receiver fibroblasts within bi-phasic GFP hydrogel taken 5 days after encapsulation. (I) Plot profile of normalized mCherry intensity distribution across the length of the hydrogel 1, 2, 3, and 5 days after encapsulation. Green shaded area indicates the region containing GFP.

Next, we attempted to activate synNotch receptors in cells embedded in 3D matrix-derived hydrogels that present synthetic ligands. To do so, we developed a click chemistry method to conjugate synthetic ligands to hydrogels (Fig. 2D). Briefly, GFP was modified with trans-Cyclooctene (TCO) NHS ester to generate GFP-TCO moieties. In parallel, gelatin polymer was modified with methacrylate (MA) groups for photo-cross-linking and methyltetrazine (mTz) to generate GelMA-mTz (Fig. S2A-B). These coordinated substitutions enable facile conjugation of TCO-modified protein ligands to the mTz-modified hydrogel polymer backbone via rapid click reaction after mixing^39, 40^ (Fig. S2C). Combining GelMA-mTz with GFP-TCO generated GelMA- GFP, which could then be photocrosslinked into a hydrogel that demonstrated retention of the GFP ligand for over seven days (S2D-E).

We then embedded anti-GFP receiver fibroblasts in GelMA-GFP hydrogels via photocrosslinking. As shown in Figure 2E-F, mCherry expression in receiver cells significantly increased in GelMA-GFP hydrogels but not in unmodified GelMA hydrogels (see also Fig. S2F- H and J). Up to 70% of the receiver cell population within GelMA-GFP had sustained activation for up to 7 days. In contrast, when we attempted to activate synNotch receiver cells via sender cells co-embedded in a GelMA hydrogel, only 30% activation of the receiver cell population was observed (Fig. S3A-B). To demonstrate spatial confinement of activation, we next encapsulated receiver fibroblasts via manual pipetting in a biphasic GelMA hydrogel, where only half of the hydrogel contained GFP. Due to the covalent linkage between GFP and GelMA, the spatial position of GFP was maintained over time and the GFP did not diffuse through the hydrogel (Fig. S2I). As shown in Figure 2G-I, mCherry activation was similarly spatially restricted to the GelMA-GFP region over time, demonstrating that the GFP ligand conjugated to the hydrogel activated synNotch only in the regions where it was originally positioned. To validate the modularity of this method, we also engineered fibrinogen-mCherry constructs via a similar click chemistry approach (Fig. S3C). We then embedded anti-mCherry/Gal4 synNotch receiver cells that activate BFP in these hydrogels. As shown in Fig. S3D, receiver cells were activated only in fibrinogen-mCherry hydrogels but not unmodified fibrinogen hydrogels. Collectively, these results demonstrate that matrix-derived hydrogels can be covalently conjugated with synthetic ligands to generate 2D or 3D materials capable of locally activating receiver cells.

### Spatial activation of synNotch via microcontact printing

Our next goal was to dictate synNotch activation patterns within multicellular tissue constructs at a spatial resolution similar to the cellular length scale. To achieve this, we adapted microcontact printing techniques designed to transfer microscale patterns of proteins (classically extracellular matrix proteins) onto culture surfaces^41, 42^. Our goal was to microcontact print GFP onto uniformly cell-adhesive surfaces (Fig. 3A). To achieve this, we treated PDMS-coated coverslips with APTES and glutaraldehyde to induce covalent bonding of proteins^43^ and then coated the surface with fibronectin for uniform cell adhesion. To optimize the transfer of GFP onto the fibronectin layer, we created simple, featureless PDMS stamps by cutting cylinders from PDMS using a biopsy punch. We coated and incubated these stamps with 0-200 ug/mL GFP solutions and then inverted them onto fibronectin-coated coverslips. Finally, we seeded coverslips with receiver cells expressing anti-GFP/tTA synNotch receptors that activate an mCherry reporter. These cells formed a confluent monolayer and demonstrated a GFP dose-dependent increase in mCherry fluorescence that saturated at roughly 100 ug/mL GFP (Fig. S4A-B), indicating that surfaces dual-functionalized with fibronectin and GFP maintained cell adhesion and activated synNotch.

**Figure 3.**
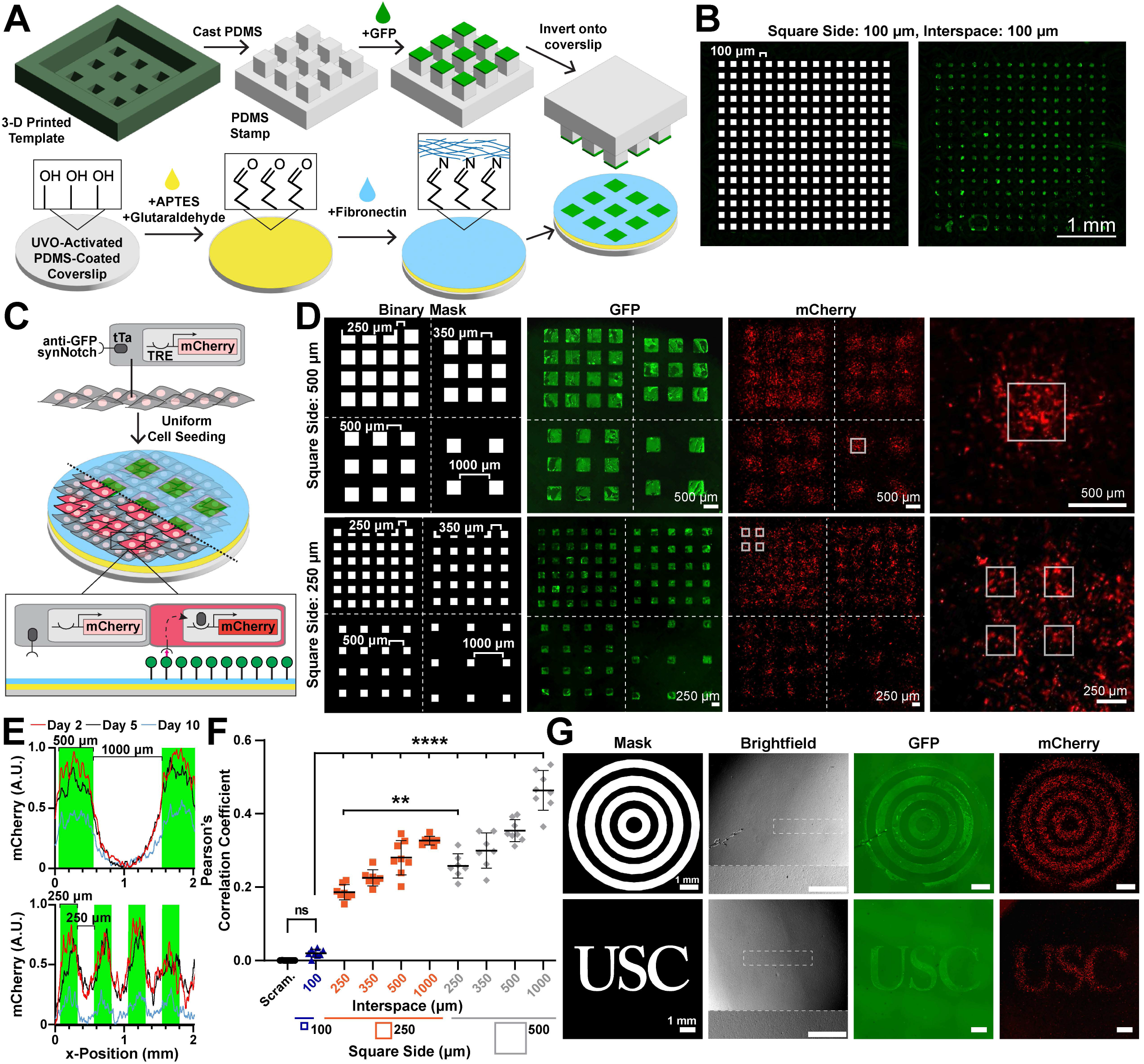
Microcontact printed GFP patterns spatially activate mCherry reporter via synNotch pathway activation. (A) Visual schematic showing the process of stamp preparation and concurrent coverslip preparation for the microcontact printing of GFP patterns. (B) Binary mask of the stamp features that contains 100µm square features with 100µm gaps and resulting GFP fluorescence image following microcontact printing (C) Schematic of anti-GFP/tTA synNotch receiver fibroblasts seeded onto GFP-patterned substrate to demonstrate local activation based on the presence of GFP. Portions of the cells were made transparent to visualize the underlying pattern of GFP. (D) Binary masks and fluorescence microscopy images of 500µm and 250µm side square GFP patterns, separated by 250, 350, 500, and 1000µm gaps, followed by mCherry fluorescence images of anti-GFP synNotch receiver cells two days following seeding. Dotted white lines separate regions with different interspaces and the solid white lines present the location of GFP patterns enlarged in the following panel. (E) Plot profile of normalized mCherry intensity on 2, 5, and 10 days following seeding onto 500×1000µm (top) 250x250µm (bottom) or GFP pattern (square side x interspace). Green indicates the regions containing GFP. (F) Quantifications of day 2 Pearson’s Correlation Coefficient (PCC) comparing Binary Mask with mCherry channel across all patterns. Scrambled binary mask images were compared to day 2 mCherry channels as a negative control. Data represents mean ± s.d, n=7-8, not significant p>0.05 (ns), p<0.01(**), p<0.0001(****). (G) Binary masks, brightfield microscopy images, and GFP fluorescence images of concentric circles and letter patterns of GFP, followed by mCherry fluorescence microscopy images taken two days following seeding onto the GFP pattern. Dotted white rectangles in the brightfield indicate the region of interest shown in higher magnification on the bottom of the same image. Scale bars, 1mm.

To induce activation of synNotch in small groups of cells within a multicellular tissue, we next developed an approach to microcontact print arrays of GFP squares with features ranging from 100 µm to 1 mm. PDMS stamps for microcontact printing are classically cast on silicon wafer templates fabricated by cleanroom-based photolithography^44^. However, this approach is not suitable for our feature sizes because they are large (100 um to 1 mm) relative to the height of photoresist conventionally used for photolithography (1-10 um). PDMS stamps with high feature to height ratios are susceptible to buckling and transfer of GFP outside the intended regions^44^. To overcome this, we used a digital light processing (DLP) 3-D printer to rapidly print templates with taller features in a photocrosslinkable resin. We first 3-D printed a template comprising an array of 100 um sided-squares with 100 um interspaces, which is roughly the resolution limit of the 3-D printer. The height of the features was set as 100 um to minimize buckling. As shown in Figure 3B, PDMS stamps fabricated in this way could successfully transfer GFP onto covalently coated FN coverslips in the intended 100 um x 100 um pattern, demonstrating successful microcontact printing using PDMS stamps cast on 3-D printed templates.

We next used these techniques to fabricate stamps and microcontact print arrays of GFP squares with sides ranging from 250 um to 1000 um and interspaces of 250 um or 500 um onto PDMS-coated coverslips pre-coated with fibronectin. The feature height for these stamps ranged from 100um to 500um, depending on square sizes and interspaces. Microcontact printed surfaces were then seeded with receiver cells with anti-GFP/tTA synNotch receptors that activate an mCherry reporter (Fig. 3C). After two days, mCherry expression was detected within the multicellular tissue in patterns that overlapped with the original design to different extents, depending on the pattern (Fig. 3D-E). To quantify the spatial fidelity of synNotch activation, we calculated the Pearson’s correlation coefficient between the binary pattern design and the mCherry images (Fig. 3F and S4E). As expected, the correlation coefficient was highest for tissues with the largest squares (500 um sides) and largest interspaces (1000 um). The correlation coefficient decreased as features and/or gaps decreased. However, for all tissues with square sizes and interspaces greater than 100 um (Fig. S4C), the correlation coefficient between the mCherry image and the binary pattern was significantly higher compared to the correlation coefficient between the mCherry image and a scrambled binary pattern with the same number of white pixels. The correlation also decreased with time due to weakening of reporter activation (Fig. 3E, S4D). Together, these data indicate that the minimum feature size for this approach is approximately 250 um. Based on this conclusion, we designed other arbitrary patterns with minimal feature sizes of 250 um, including concentric circles and letters. Qualitatively, we observed similar agreement between the binary pattern, GFP fluorescence, and mCherry fluorescence (Fig. 3G), demonstrating versatility of pattern designs.

Our next goal was to scale-up this approach to spatially activate multiple distinct genetic programs in the same multicellular tissue. Previous studies have demonstrated that two synNotch receptors can be integrated into a single dual-receiver cell^23^. Thus, we asked if culturing dual-receiver cells on a surface patterned with two synthetic ligands in distinct arrangements would generate a tissue with corresponding patterns of distinct genetic programs (Fig. 4A). We first generated a dual-receiver fibroblast cell line (C3H) that harbors an anti-GFP/tTA synNotch receptor that activates an miRFP reporter and an anti-mCherry/Gal4 synNotch that activates a BFP reporter (Fig. S5C). To validate the responses to synthetic ligands of these cells, we seeded them on a culture surface microcontact printed with a uniform layer of GFP, mCherry, or both. As shown in Fig. S5A-B, miRFP was expressed only on GFP surfaces and BFP was expressed only on mCherry surfaces, demonstrating orthogonal activation of the two pathways. On surfaces with both GFP and mCherry, both miRFP and BFP were expressed, indicating activation of both pathways. Next, to prototype the generation of spatial patterns of gene expression starting from a uniform population of dual-receiver cells, we adsorbed GFP from a droplet in one corner of a culture surface and a droplet of mCherry in the opposing corner. Dual-receiver cells cultured uniformly on the surface activated miRFP and BFP in a spatial pattern corresponding to the GFP and mCherry droplets, respectively, demonstrating macroscale spatial control over the activation of two synNotch pathways in one cell population (Fig. S5D-E). Finally, to provide more precise spatial control over the patterns, we microcontact printed an array of 500 µm-wide rows of GFP with 500 um interspacing. We then stamped perpendicular mCherry rows by manually positioning the orientation of the stamp (Fig. 4B).

**Figure 4.**
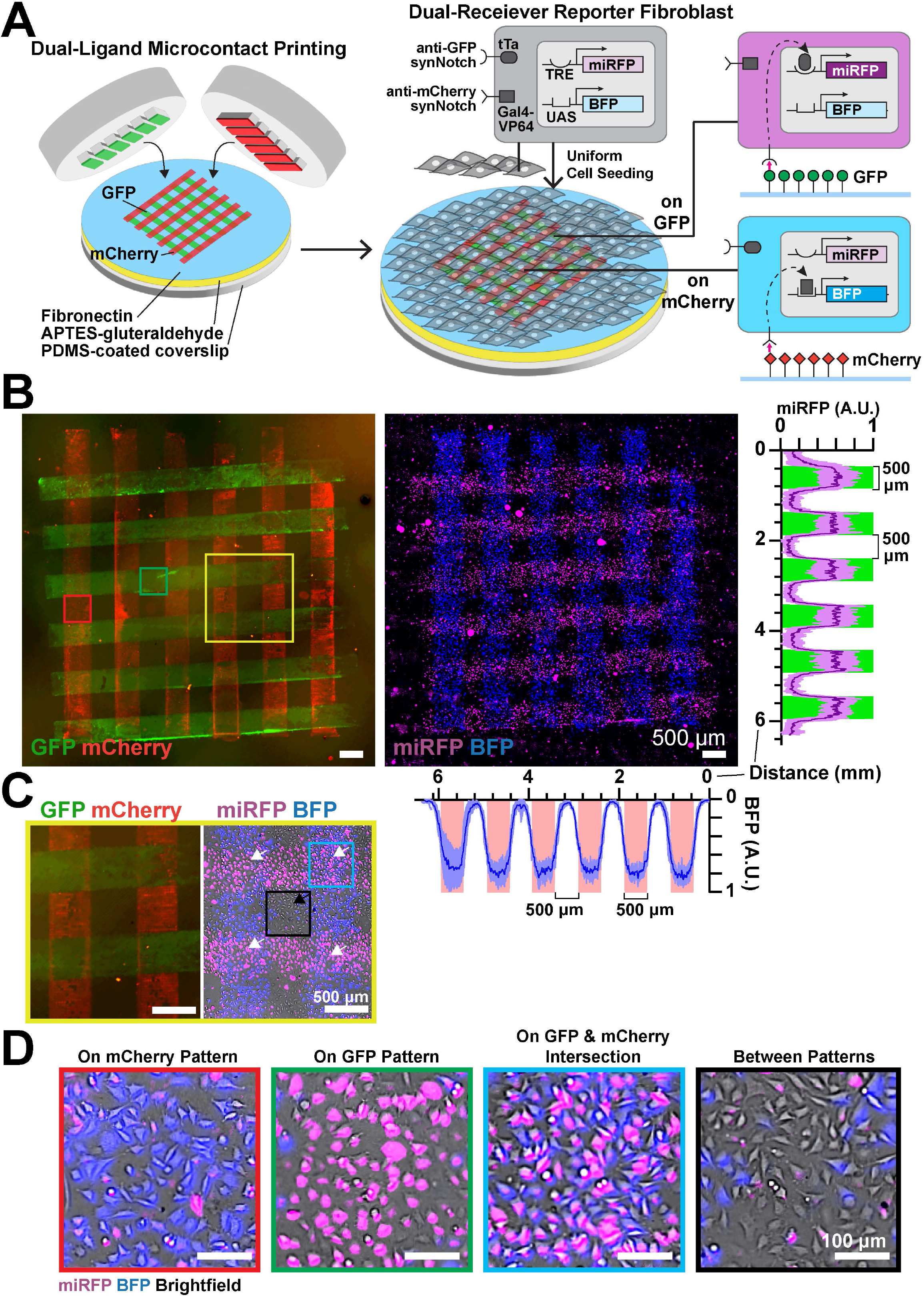
Patterned GFP and mCherry spatially activate dedicated reporter genes via synNotch activation in dual receiver fibroblasts. (A) Schematic showcasing: (left) dual-ligand microcontact printing, and (right) seeding of dual-receiver L929 cell where anti-GFP synNotch drives miRFP reporter gene and anti-mCherry synNotch orthogonally activates BFP. (B) Fluorescence microscopy images of microcontact printed GFP and mCherry perpendicular rows of 500µm width and subsequent dual reporter expression taken 24 hours after uniform seeding. Scale bars, 500μm. (Right) Normalized plot profiles of miRFP intensity across each row axis of engineered dual-receivers 24 hours following seeding onto perpendicular GFP and mCherry patterns. Green bars indicate regions containing GFP. Line profiles represent mean ± s.d, n=7. (Below) Normalized plot profiles of BFP intensity across each row axis of engineered dual- receivers 24 hours following seeding onto perpendicular GFP and mCherry patterns. Red bars indicate regions containing mCherry. Line profiles represent mean ± s.d, n=7. Regions of interest on mCherry (red border), on GFP (green border), or on intersecting patterns (yellow border) enlarged in the following panels. (C) Higher magnification fluorescence microscopy images of GFP and mCherry perpendicular rows and subsequent dual reporter expression/brightfield image. White arrows indicate miRFP+ and BFP+ double positive cells in regions where GFP/mCherry overlap. Black arrows indicate double negative cells in regions where no ligands were patterned. Regions of interest on GFP and mCherry intersection (blue border) or non-patterned region (black border) enlarged in the following panels. Scale bars, 500μm. (D) Higher magnification fluorescence microscopy images to demonstrate BFP and miRFP expression by dual-receiver fibroblasts on mCherry, GFP, intersection, and between patterns. Scale bars, 100μm.

When seeded with dual-receiver cells, we observed rows of miRFP-expressing cells perpendicular to rows of BFP-expressing cells (Fig. 4B-D and Fig. S5F), as expected. At the GFP and mCherry intersections, cells expressed both miRFP and BFP, indicating activation of both synNotch pathways (Fig. S5G), generating 4 reporter “states” for the initially uniform population of engineered cells (parental, BFP+, miRFP+, BFP+/miRFP+ double positive) within the 1.5mm^2 tissue. Thus, two independent synNotch genetic programs can be spatially controlled by culturing dual-receiver cells on user-defined patterns of the two synthetic ligands, to generate a multicellular tissue with up to 4 spatially controlled reporter gene expression states.

### Spatial control of concurrent differentiation to skeletal muscle and endothelial cell precursors

Beyond expression of fluorescent reporter proteins, synNotch receptors have also been used to activate transgenes that control cell phenotypes or behaviors via overexpression of transcription factors, such as Snail for epithelial to mesenchymal transitions or myoD for transdifferentiation of fibroblasts to skeletal muscle precursors^23^. Thus, we next tested if synthetic ligands presented by materials could drive overexpression of functional transcription factors that induce transdifferentiation.

We first generated fibroblasts expressing an anti-GFP/tTA synNotch receptor that activates myoD (Fig. 5C). Previous studies have shown that micromolded gelatin hydrogels are favorable for myotube adhesion and alignment^33, 45^. Thus, to test if these surfaces could be used to transdifferentiate and align synNotch-induced myotubes, we constructed gelatin hydrogels that are either featureless or micromolded with 10 um ridges separated by 10 um spacing and then enzymatically conjugated GFP to the surface using the procedure described above (see Fig. 2A). When cultured on these surfaces, anti-GFP synNotch receiver cells induced myoD and fused into alpha-actinin-positive myotubes aligned with the micromolded ridges (Fig. 5A-B and S6A).

**Figure 5.**
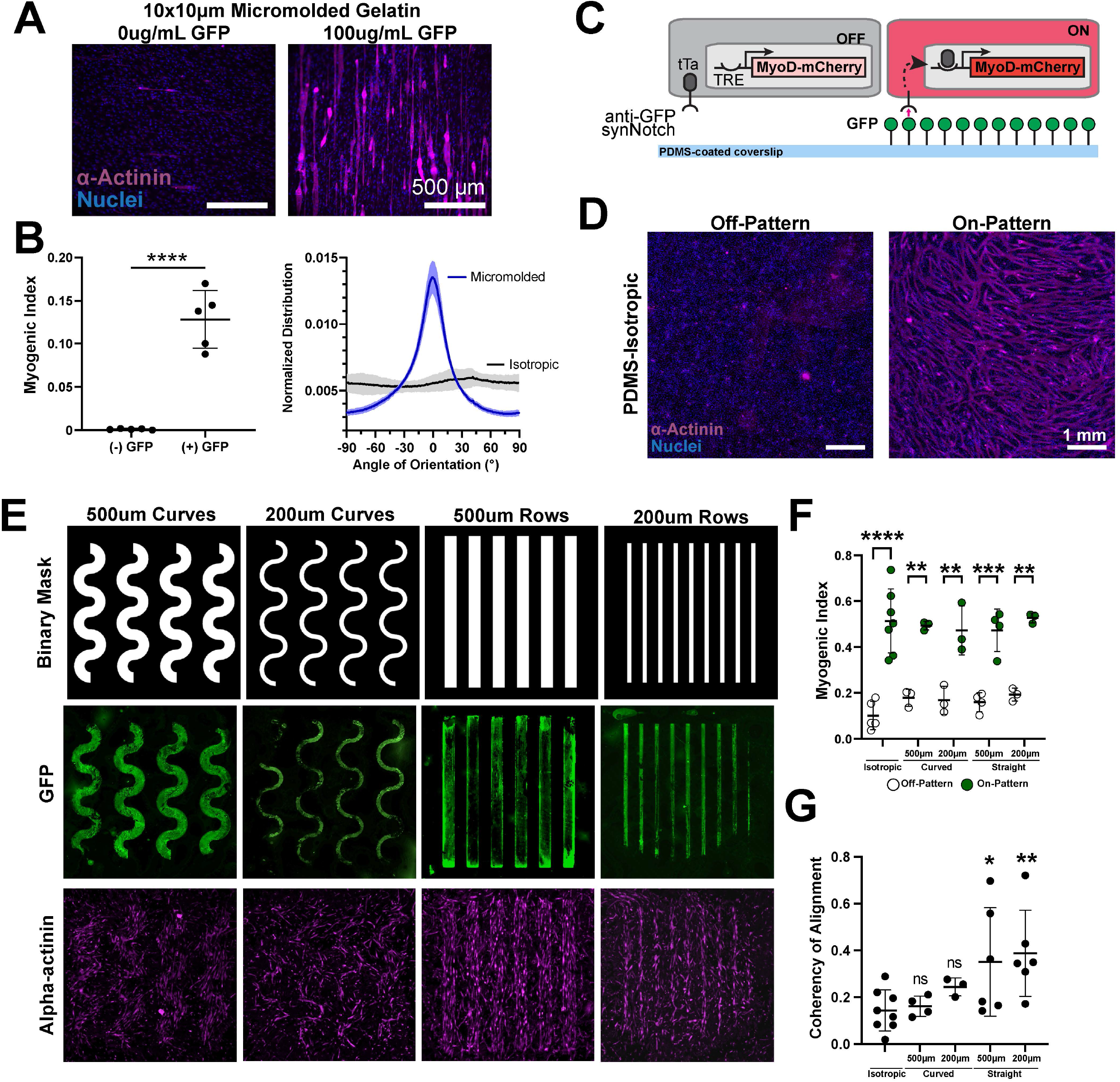
Microcontact printed GFP patterns spatially activate myoD and initiate myotube differentiation in embryonic fibroblasts via synNotch. (A) Fluorescence images of anti-GFP/tTA synNotch receiver fibroblasts stained for α-actinin 7 days following seeding onto 10×10µm grooves micromolded gelatin substrates with 0 or 100µg/mL transglutaminase-conjugated GFP. Scale bars, 500μm. (B) Myogenic index in the presence or absence of 100µg/mL transglutaminase-conjugated GFP on 10×10µm grooves micromolded gelatin, quantified by image analysis. Data represents mean ± s.d, n=5, p<0.0001(****). Myotube alignment of α-actinin stained myotubes on GFP-conjugated 10×10µm groove micromolded gelatin compared to isotropic (no topography) gelatin substrate, quantified by image analysis. Line plot represents average angles of orientation distribution of 5 individual images from an individual sample. Data represents mean ± s.d. (C) Schematic of embryonic fibroblasts expressing anti-GFP synNotch activating myoD and mCherry transgenes seeded onto GFP patterned PDMS substrate. (D) Sarcomeric α-actinin staining on isotropic GFP patterns on a PDMS substrate, stained three days following seeding on GFP patterns. Off-Pattern indicates image taken on GFP-negative regions, On-Pattern indicates images taken within GFP-positive regions. Scale bars, 1mm. (E) Binary mask used to generate stamps for microcontact printing followed by fluorescence microscopy images of GFP ligand and subsequent α-actinin staining for each pattern: 500µm curves, 200µm curves, 500µm rows, and 200µm rows. Samples were stained three days following uniform seeding onto GFP patterns. (F) Day 3 Myogenic Index, quantified with image analysis, on and off GFP for each pattern (isotropic, 200,500 curves, 200,500 straight rows). Data represents mean ± s.d, n=3-6, p<0.01(**), p<0.001(***), p<0.0001(****). (E) Coherency of myotube alignment measured across all patterns quantified with image analysis. Significance values compared to isotropic control. Data represents mean ± s.d, n=3-8, p<0.05(*), p<0.01(**).

Another approach for engineering aligned muscle tissues is to culture muscle cells on microcontact printed lines of matrix proteins [Feinberg Parker biomaterials]. We tested if this approach was compatible with synNotch by microcontact printing lines of a mixture of fibronectin and GFP. When the same receiver cells were cultured on these surfaces, they transdifferentiated into aligned myotubes (Fig. S6B), indicating that microcontact printing matrix proteins and synthetic ligands can be used to both control tissue architecture and transdifferentiation.

In these approaches described above, a population of fibroblasts was uniformly transdifferentiated to myoblasts. Our next goal was to selectively transdifferentiate fibroblasts to myoblasts in a spatially controlled manner as a first step towards generating tissues with multiple distinct cell types arranged in prescribed patterns. To achieve this, we wanted to use the approach described above (Fig. 3) to microcontact print rows of GFP on fibronectin-coated surfaces. To test that microcontact-printed GFP can induce transdifferentiation of anti-GFP/tTA synNotch receiver cells, we cultured these cells on surfaces uniformly microcontact printed with GFP. As shown in Fig. 5C-D, receiver cells transdifferentiated to multi-nucleated, alpha-actinin positive myotubes on the GFP surfaces. We next wanted to see if we could achieve spatially controlled differentiation only on certain regions of the culture; to test this, we printed thin or thick, curved or straight rows and then seeded the printed surfaces with fibroblasts harboring an anti-GFP synNotch receptor that activates MyoD (Fig. 5E). After three days, we fixed and stained tissues for alpha-actinin and quantified the myogenic index on and off the pattern by using the binary pattern as a mask (Fig. 5F). Myogenic index was significantly higher on-pattern compared to off-pattern for all geometries, demonstrating local geometric control of transdifferentiation. We also quantified the coherency of the tissues as a proxy for alignment and observed higher coherency for tissues on the straight rows compared to the curved rows (Fig. 5G see also Fig. S6C-G). Thus, we can selectively transdifferentiate fibroblasts to myoblasts in a geometrically prescribed way while also controlling the global alignment of the tissue, demonstrating that we can separately and concurrently control local differentiation and tissue architecture.

To exploit the modularity of this technology, we next tested if transdifferentiation to another cell fate could be activated by a similar approach. Due to the universal need for vascularization in engineered tissue constructs, including muscle, we focused on transdifferentiating fibroblasts to endothelial cells precursors, which was previously shown via doxycycline-inducible overexpression of the master transcription factors ETV2^46, 47^. Thus, we generated fibroblast receiver cells engineered with an anti-mCherry/Gal4 synNotch receptor that activates an ETV2- BFP cassette (Fig. 6A). We then passively adsorbed mCherry onto culture surfaces, cultured ETV2 receiver cells on them for three days, and then fixed and stained the cells for endothelial cell precursor markers. As shown in Fig. 6B-C (see also Fig. S7A-E), the fibroblasts transdifferentiated to VEGFR2-positive endothelial precursors that also expressed VE-cadherin on their membrane, demonstrating that we engineered a receiver fibroblast cell line that can transdifferentiate to endothelial cell precursors via mCherry synthetic ligands presented on a plate. To test if we can control the geometry of transdifferentiation in this context too, we generated uniformly adhesive surfaces and then microcontact printed mCherry as user-defined row patterns of different widths and interspaces. As shown in Fig. 6D-E, when anti-GFP synNotch cells that induce ETV2-BFP were cultured on these surfaces, the transgene was induced only locally on the patterns at day 1 and day 3. We then designed a pattern to replicate a branching network structure typical of vascular beds^48^ and showed the formation of a tissue consisting of activated cells in the corresponding pattern surrounded by a uniform layer of fibroblasts (Fig. 6F and S7D). Thus, microcontact printed ligands can activate SynNotch-induced transdifferentiation to skeletal muscle precursors and endothelial precursors with spatial control.

**Figure 6.**
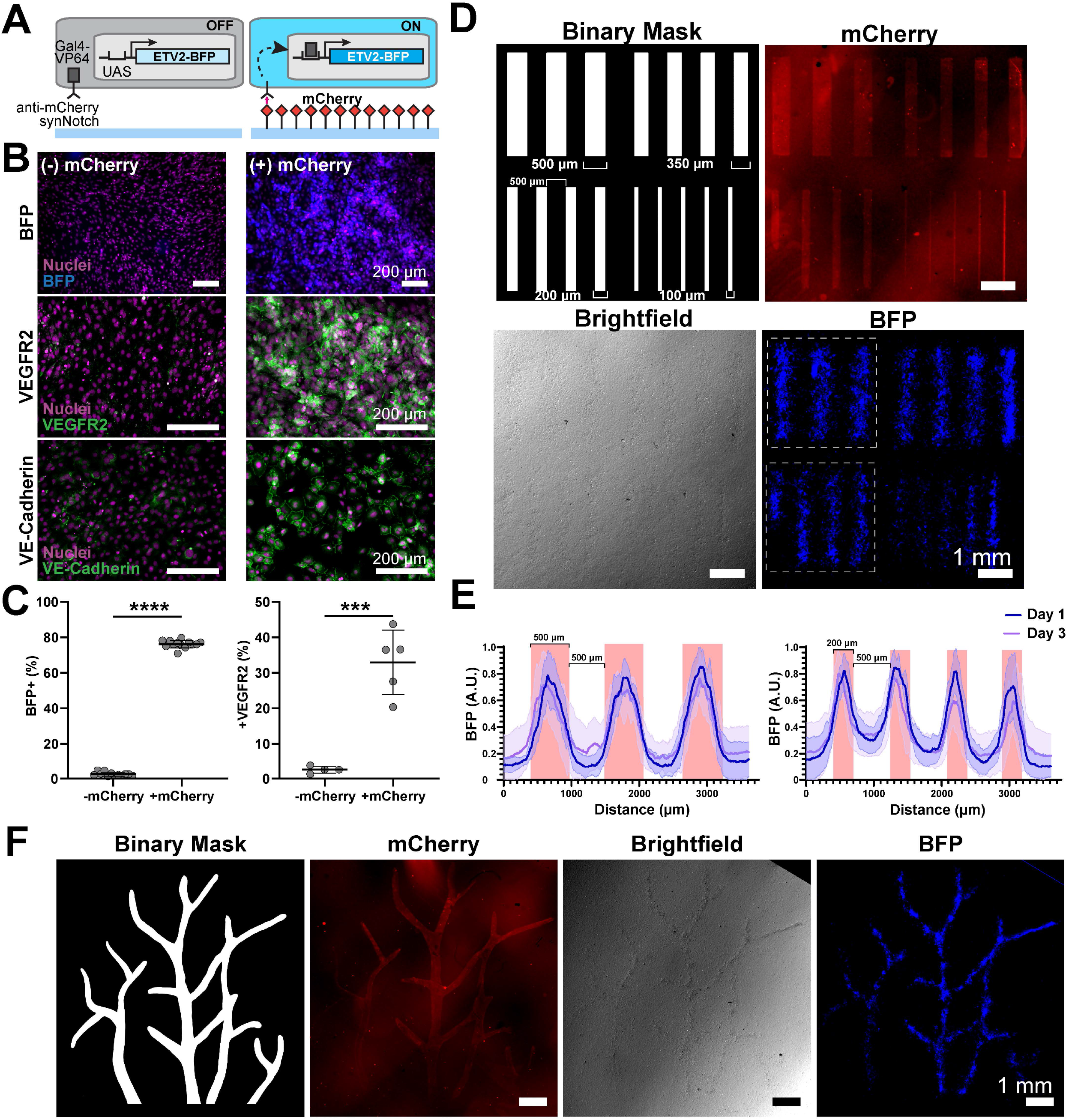
mCherry activates ETV2 and reporter BFP to induce endothelial differentiation in embryonic fibroblasts via synNotch. (A) Schematic of embryonic fibroblasts cell line (C3H) expressing anti-mCherry/Gal4 synNotch activating ETV2 and BFP transgenes seeded onto mCherry patterned substrate. (B) Fluorescence microscopy images of BFP reporter (top), endothelial markers VEGFR2 (middle) and VE-Cadherin (bottom) stained 3 days following seeding onto control wells (-mCherry) or plate- dried mCherry (+mCherry). Scale bars, 200μm. (C) Percent of cells expressing BFP (left) and VEGFR2 (right) in the presence (+mCherry) or absence (-mCherry) of mCherry, quantified with flow cytometry. Data represents mean ± s.d, BFP n=12-13, VEGFR2 n=4-5, p<0.001(***), p<0.0001(****). (D) Binary mask used to generate stamps for microcontact printing followed by fluorescence microscopy images of Day 0 mCherry pattern, brightfield, and Day 1 BFP expression. Scale bars are 1mm. White dotted squares designate representative fields used for quantification in E. (E) Plot profile of normalized BFP intensity on Day 1 and Day 3 following seeding onto 500µm (left) and 200µm (right) mCherry rows within a row matrix. Red bars indicate the regions containing mCherry. Line profiles represent mean ± s.d, n=6. (F) Binary mask used to generate stamps for microcontact printing followed by fluorescence microscopy images of Day 0 mCherry pattern, brightfield, and Day 1 BFP expression for vasculature-like pattern. Scale bars are 1mm.

Finally, we asked if we could engineer a tissue construct in which multiple distinct cell fates are arranged in user-specified geometries. To do so, we first engineered a “dual-lineage” cell line with two synNotch pathways: an anti-GFP/tTA synNotch receptor that activates myoD-miRFP and an anti-mCherry/Gal4 synNotch that activates ETV2-BFP (Fig. 7A center). To test the functionality and orthogonality of these pathways, we cultured these cells on surfaces with a uniform coating of GFP or mCherry for three days and then stained them for markers of differentiation. As shown in Fig. S8A-C, these cells transdifferentiated to alpha-actinin-positive muscle precursor cells or VEGFR2-positive endothelial precursor cells, respectively. As a curiosity, we evaluated the effects of culturing cells on both ligands, which would induce overexpression of both myoD and ETV-2 in the same cells. In this case, it seemed that transdifferentiation to both pathways was impaired, as these cells did not appear to differentiate towards skeletal muscle nor express endothelial cell markers (Fig. S9E). To prototype simple spatial activation, we used a micropipette to deposit droplets of GFP and mCherry in opposing corners of a culture surface (Fig. S9A-B). Dual lineage cells cultured on this surface activated the fluorescent protein reporters with expected spatial control, and displayed multinucleation in the GFP-coated region, indicating feasibility for spatial activation of differentiation.

**Figure 7.**
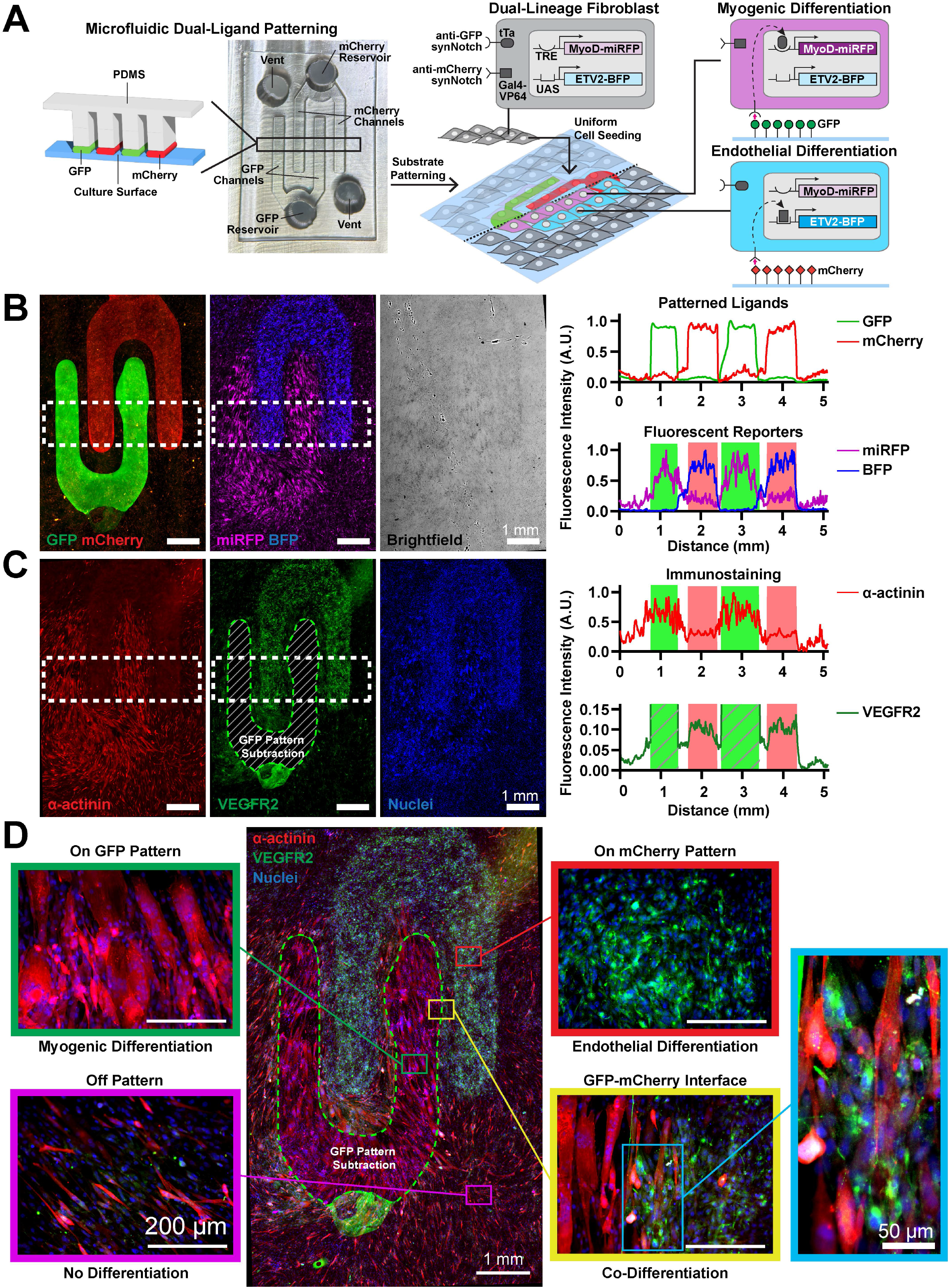
Spatially controlled co-transdifferentiation of myogenic and endothelial fates via dual-lineage synNotch engineered cell and dual ligand adsorption in microfluidic device. (A) Schematic showing dual protein patterning technique using capillary-driven microfluidic patterning, based on shallow and deep channels, to generate parallel lines of GFP and mCherry. Feature size, 500 µm. Schematic of dual-lineage mouse embryonic fibroblasts (C3H line) expressing anti-GFP/tTA synNotch that activates MyoD and miRFP as well as an anti-mCherry/Gal4 synNotch that orthogonally activates ETV2 and BFP transgenes, seeded onto GFP and mCherry patterned substrate. (B) Fluorescence images of GFP and mCherry ligand (left), reporter genes expression (center) and brightfield (right) of dual-lineage cells 3 days following uniform seeding onto capillary-fluidic ligand patterns. Dotted white rectangles represent regions of interest for quantification. Scale bars, 1mm. Plot profiles on the right show normalized fluorescence intensity of ligands and reporters taken in the region of interest. Green and red bars indicate regions containing GFP or mCherry, respectively. (C) Fluorescence images of immunostained α-actinin (left), VEGFR2 (center), and nuclei (right), stained 3 days after uniform seeding onto ligand patterns. Dotted white rectangles represent regions of interest for quantification; region with striped lines, delimited by a dotted green line, represents a region where the GFP signal has been subtracted to visualize VEGFR2 staining. Scale bars, 1mm. On the right, plot profiles of normalized fluorescence intensity of α-Actinin and VEGFR2 taken across the region of interest. Green and red bars indicate regions containing GFP or mCherry respectively. Striped green bars indicate regions where the GFP signal has been subtracted (see Fig. S9D for subtraction procedure). (D) Center, merged fluorescence image of α-Actinin and VEGFR2 immunostaining on dual-lineage cells uniformly seeded onto GFP and mCherry pattern. Dotted green lines represent region where the GFP signal has been subtracted to visualize VEGFR2 staining. Scale bar, 1mm. Around the central image, higher magnification fluorescence images taken within distinct regions of the pattern are shown: on mCherry (red border), interface between mCherry and GFP regions (yellow border), on GFP (green border), and off pattern (purple border). Scale bar, 200µm. Higher magnification of interface (blue border), visualizing α- Actinin and VEGFR2 staining is shown on the far right with scale bar of 50µm.

Next, we wanted to pattern multiple synthetic ligands onto a surface simultaneously and with spatial control. To do so, we adapted approaches for controlling the distribution of multiple streams of liquids with an open capillary microfluidic device^49^. Briefly, the intended fluid paths are created as shallow channels that are laterally open and adjacent to deep channels. Fluids preferentially travel along the shallow channels instead of the deep channels because of greater surface tension in shallow channels. We used this concept to design a microfluidic device for delivering solutions of GFP and mCherry as interdigiting 500 µm wide rows (Fig. 7A - left). We printed inverse templates of the device using a DLP 3-D printer, which were then cast with PDMS. Air vents and GFP and mCherry reservoirs were punched into the PDMS and the device was attached to a culture surface and loaded with GFP and mCherry solutions. After overnight incubation, the PDMS was removed and remaining solutions were briefly air dried, leaving behind interdigitating rows of GFP and mCherry adsorbed on the surface (Fig. 7B). When dual-lineage cells were cultured on these surfaces, cells adhered uniformly to the entire surface and proceeded to transdifferentiate to myoblasts or endothelial cells in a pattern corresponding to the intended pattern of ligands (Fig. 7B-C). As highlighted in Fig. 7D, alpha-actinin-positive muscle precursor cells were confined to the GFP rows, VEGFR2-positive endothelial precursor cells were confined to the mCherry rows, and intermixing of these two cell types was observed at the interface between GFP and mCherry. Cells on the unpatterned regions remained fibroblasts. Of note, these tissues were maintained in standard cell culture media, without the need for soluble differentiation factors or biophysical stimulation to drive cell fates. Thus, by integrating microfabrication techniques with synNotch receptors, we induced a uniform population of fibroblasts to differentiate into a tissue with three distinct cell populations (skeletal muscle precursors, endothelial precursors, and fibroblasts) patterned in user-defined microscale geometries.

## DISCUSSION

In this study, we engineered several material-to-cell signaling pathways to spatially activate user-defined genetic programs in multicellular systems. We achieved this by engineering cells with synNotch receptors to define cellular inputs and outputs while concurrently engineering materials to present synthetic ligands with different ranges of spatial control. The variety of materials for synthetic ligand presentation yields powerful and highly flexible tools for activating material-to cell pathways. Due to the functional modularity of synNotch receptors, material-activated pathways can be used to drive any number of transcriptional programs or differentiation pathways. Thus, these generalizable technologies provide users with unprecedented abilities to dictate spatial patterning of gene expression in multicellular constructs.

Because of the highly powerful level of transcriptional control over cell behaviors in natural systems, many efforts in synthetic biology have focused on engineering sophisticated transcriptional circuits^50, 51^. In the area of stem cell and cell differentiation, genetic overexpression of master transcription factors has demonstrated robust control over cell differentiation^52, 53^. The capacity to induce master transcription factors with user-defined spatial control has inspired recent advances, such as engineering cells with light-activatable signaling pathways to gain spatiotemporal control over cell behaviors with light^54, 55^. For example, myoD overexpression has been induced by a genetically integrated optogenetic switch that can be activated spatially^56^. Although optogenetic approaches have the potential for powerful spatiotemporal control over cell behaviors, optogenetic technologies require sophisticated light manipulation devices and cannot control multiple cell behaviors concurrently. With our previous development of synNotch, we generated a novel way to activate user-defined genetic programs via user-defined ligands presented by neighboring cells. Here, we advanced this technology to a new level by activating synNotch via multi-modal materials commonly used for tissue engineering. Importantly, we showed that this novel approach can be used to define spatial patterns of not only gene expression, but also differentiation.

To present synthetic ligands, we modified several different types of materials, each with trade-offs. By engineering cells to secrete fusions of synthetic ligands and natural matrix proteins (e.g., fibronectin), ligands are presented in a natural extracellular matrix network comprising a diversity of endogenous macromolecules, which may enhance receiver cell adhesion and survival. However, spatial control is very coarse, as cells inherently migrate and deposit matrix fibers semi-stochastically. Hydrogels are the most common class of materials for tissue engineering due to their high water content and multiple tunable properties, including stiffness, porosity, and composition^57^. Thus, we developed versatile and modular methods for presenting synthetic ligands via hydrogels, including: conjugation of ligands on the surface of hydrogels for 2-D culture or in the bulk of hydrogels for 3-D culture; conjugation by an enzymatic reaction or click chemistry reaction; conjugation of GFP or mCherry as synthetic ligands; and conjugation of ligands to hydrogels with gelatin or fibrinogen polymer backbones. One limitation is that spatial patterning of ligands in hydrogels was limited to manual pipetting, which is relatively coarse. However, many other modalities exist to conjugate and/or release proteins from hydrogels with spatial control^58^, which can be integrated in future iterations.

To achieve greater spatial control, we microfabricated PDMS stamps and microfluidic devices to pattern synthetic ligands onto 2-D surfaces. By fabricating these components on 3-D printed templates instead of classical photolithography-based wafers, we also achieved a wider range of pattern designs and more rapid prototyping capabilities, with the tradeoff that spatial resolution was confined to 100 um or above. To pattern synthetic ligands at sub-cellular spatial resolution, photolithography would still be required. Our two PDMS-based technologies also have unique tradeoffs. Microcontact printing can be used to generate essentially any geometrical pattern (including isolated islands) but cannot be used to precisely position multiple ligands since each stamp must be positioned manually. Conversely, registering the placement of multiple ligands is possible with a microfluidic device, but pattern geometries are limited to continuous channels connected to a reservoir. Thus, these constraints must be considered when choosing a patterning modality.

Our most sophisticated tissue construct comprised interdigitating rows of skeletal muscle and endothelial cells, with some intermingling of the cells at the interface. Importantly, the skeletal muscle cells and endothelial cells were co-transdifferentiated from a uniform population of fibroblasts. This approach is in contrast to conventional tissue engineering techniques, which generally differentiate individual cell types in isolation and then combine them. Our approach may better mimic natural tissue morphogenesis, where multiple cell fates emerge simultaneously from a uniform cell population. Studies have also shown that supporting cells, such as endothelial cells, improve the maturation of human induced pluripotent stem cell- derived cardiomyocytes^59, 60^, and that co-differentiation of different lineages concurrently more closely recapitulate the conditions occurring during embryonic development^61^. An interesting hypothesis to explore with our technology is whether co-differentiation of supporting cells (e.g., endothelial cells) adjacent to parenchymal cells (e.g., muscle cells) has additional benefits for phenotypic maturity.

One interesting observation from our data is that different ligand-presenting materials yielded different temporal patterns of synNotch activation. For example, synNotch activation peaked at three days and then subsided when ligands were microcontact printed on PDMS, whereas activation was more sustained when synNotch was activated by ligands conjugated to 3-D hydrogels. This could be caused by differences in the conjugation of the ligands to the materials, such as the strength of the material-ligand bond or ligand orientation, and/or differences in the activation of the synNotch receptor itself. The mechanism of transduction by synNotch receptors is thought to proceed similarly to endogenous Notch receptors. In the core regulatory region of endogenous Notch receptors, a pulling force is generated upon ligand binding, which exposes a protease cleavage site for a protease that is constitutively active in the membrane; this cleavage then liberates the intracellular domain which is a transcription co- activator^26, 62^. The mechanisms of activation of Notch and synNotch receptors via cell-presented ligands have been compared, individuating possible mechanisms of activation that distinguish different synthetic and natural receptor constructs^63^. Thus, the mechanism of activation of synNotch by material-presented ligands may also differ from cell-presented ligands and may differ for different materials. Increased mechanistic understanding of synNotch activation by materials could yield increased capacity for spatial, and perhaps temporal, control of gene expression.

We anticipate that our novel approach for activating synthetic pathways for transdifferentiation by a material can be combined with other technologies for deriving complex *in vitro* tissues, such as organoids^64^. To generate organoids, stem cells are exposed to natural ligands that orchestrate their self-organization into complex cellular arrangements^64^. However, although cellular complexity at the microscale in organoids is remarkably similar to endogenous organs, users lack geometric control over the arrangement of cells at higher levels, leading to tissue constructs that are largely heterogeneous and poorly reproducible with un-natural architectural features. Combining synthetic biology and organoids is a recognized frontier of the field^65–69^ and synNotch-mediated spatial patterning technologies, such as those presented here, could represent a step in the direction of ultimate user-control of cell behaviors across multiple spatial scales for engineering *in vitro* multicellular systems.

## Contributions

M.G., T.H., T.M., S.D., A.R.M., N.C., S.L., M.M., L.M. designed the experiments. M.G., T.H., T.M., S.D., N.P., R.E.L., J.S., B.J., performed the experiments. M.G., T.H., T.M., S.D., R.E.L., B.J., analyzed the data. M.G., T.H., T.M., S.D., A.K., S.L., M.M., L.M. contributed to data interpretation and discussion. M.G., T.H., T.M., S.D., S.L., M.M., L.M. wrote the manuscript.

## Acknowledgments

The authors acknowledge Marion Johnson all the members of the Morsut, McCain, Li, Khademhosseini Lab for insightful discussions and suggestions on the project. The authors acknowledge their family and friends that support them always and in particular for the times of this work that took place during the COVID-19 pandemic. This project was supported by NSF RECODE from CBET-2034495 (MLM, LM, R35 from NIGMS R35 GM138256 (LM), USC Department of Stem Cell Biology and Regenerative Medicine Startup Fund (LM), Viterbi Center for CIEBOrg (LM MLM), TH acknowledges support from the “Ruth L. Kirschstein National Research Service Award T32HL069766 and the UCLA Eli and Edythe Broad Center of Regenerative Medicine and Stem Cell Research, Research Award. Fellowships for students: CIRM fellowship for SD, Fellowships from BME Department for first year for MG, NC and SD. Grace True, Finacy Jin for technical support for the project.

## METHODS

### Genetic Constructs Design

Fibronectin-GFP plasmids were generated from PiggyBac backbone and FN-YPET (Addgene #65421). GFP and mCherry-responsive synNotch construction: pHR_SFFV_myc- LaG17_synNotch_TetRVP64 (Addgene plasmid# 79128) and pHR_EF1a_flag- LaM4_synNotch_Gal4-VP64, built from pHR_EF1a_flag-LaM4_synNotch_TetRVP64 (Addgene plasmid#162237) and HR_pGK_LaG17_synNotch_Gal4VP64 (Addgene plasmid# 79127). The response-element plasmids pHR_TRE_MyoD-P2A-mCherry, pHR_TRE_MyoD-P2A- miRFP703_PGK_PuromycinR, and pHR_UAS_ETV2-P2A-tBFP_PGK_HygromycinR (with and without transcription factor) were generated from pHR_TRE, pHR_5x Gal4 UAS (Addgene plasmid# 79119), mouse MyoD (NP_034996.2), and mouse ETV2 (NP_031985.2). All constructs were cloned via In-Fusion HD Cloning (Takara Bio).

### Lentivirus Production

Lentivirus was produced by cotransfecting pHR clones plasmids with vectors encoding packaging proteins (psPAX2, pVSVG) using Lipofectamine LTX (ThermoFisher) into 70-80% confluent HEK- 293T cells within 6-well plates. Viral supernatants were collected 2-3 days after transfection, sterile filtered with 0.45μm PES (Genesee Scientific), and used directly or 10x concentrated using LentiX Concentrator (Takara Bio) following manufacturers instructions prior to adding to cell lines.

### Cell Culture

L929 mouse fibroblast cells (ATCC# CCL-1), HEK293 cells (Takara 632180), C3H/10T1/2 Clone 8 (ATCC# CCL-226), and NIH/3T3 (ATCC# CRL-1658) were cultured in DMEM (ThermoFisher) supplemented with 10% Fetal Bovine Serum (ThermoFisher) and 100U/mL penicillin/streptomycin (ThermoFisher). Cultures were maintained in an 37°C incubator with 5% CO2 and relative humidity (VWR).

### Cell Line Engineering

2×10^4^ 3T3 cells were seeded in a 12-well plate. The following day cells were transfected with 1 ug FN-eGFP PiggyBac plasmid using 2.5 μL Lipofectamine LTX with 1 μL Plus reagent diluted in 100 μL OptiMEM. Transfected cells were selected using 2 μg/mL Puromycin. Additionally, the established line was transduced with lentivirus encoding the expression of constitutive H2B- miRFP703.

For viral transduction, 20-50μL concentrated (or equivalent non-concentrated) viral supernatant(s) were added to 5-10×10^4^ suspended cells supplemented with 10μg/mL polybrene (Sigma), then transferred into a 12-well plate for 2-3 days before changing to fresh media. Following transduction, all applicable cell lines were selected using Puromycin (L929 – 10μg/mL, C3H – 1 ug/mL, NIH3T3 2 μg/mL, ThermoFisher) and Hygromycin B (L929, C3H – 400μg/mL, MedChem Express) for the expression of transgenes. Cells were sorted for the coexpression of each component via fluorescence-activated cell sorting on a FACS ARIA II (Beckton-Dickinson) by staining with appropriate fluorescently tagged anti-Myc and anti-Flag antibody for 30 minutes at 4°C (Cell Signaling Technologies) or expression of the transgenes. A bulk-sorted polyclonal population of engineered cells were used for experiments, unless otherwise noted. For single-cell clonal populations, single cells were sorted individually into 96-well plates from selected and stained populations using a FACS ARIA II.

### GFP and mCherry Production

GFP, mCherry, and GFP-LACE (pET28-His6-GFP-C-LACE, gift from Jeffrey Bode Addgene plasmid # 133913) were purified as an N-terminal hexahistidine fusion protein. To express GFP, BL21(T1R) E. coli cells were grown to an optical density of 0.5 from an overnight-grown glycerol stock, chilled to 25°C, induced with 1 mM IPTG and allowed to express for 5 hours. To express mCherry, BL21-AI E.coli (Thermo Fisher) were transformed with mCherry-pBAD (gift from Michael Davidson & Nathan Shaner & Roger Tsien, Addgene plasmid # 54630), grown to an optical density of 0.6 from an overnight-grown glycerol stock, induced with 0.04% w/v L-Arabinose (Sigma), and allowed to express for 5 hours, based on previous studies (https://dergipark.org.tr/en/pub/iarej/issue/44303/429547). The proteins were purified by NEBExpress Ni Spin Columns (New England Biolabs) following manufacturer’s instructions, dialyzed against 1x PBS overnight at 4°C, sterile filtered, and frozen at -80°C until use.

### Microparticle-conjugation and Activation

Carboxylated magnetic polystyrene microparticles (Magsphere, MCA5UM) were first washed in 0.1M MES (pH 5.8), activated with 250mM 1-Ethyl-3-(3-dimethylaminopropyl)carbodiimide (EDC, Sigma)/ N-Hydroxysuccinimide (NHS, Sigma) in MES for 15 minutes, then washed two times with PBS (pH 7.4). Particles were then incubated in varying concentrations of GFP in PBS (0- 1000μg/mL) overnight at 4°C with inversion mixing. All washing steps used EasySep Magnet (Stemcell Technologies) or centrifugation (8000g for 5 minutes) to change solutions. Particle concentration was determined using hemocytometer and directly loaded into cell suspensions (10 particles per cell) prior to seeding L929 anti-GFP synNotch receiver cells 25,000 cells/cm^2^ directly into wells or gelatin-coated coverslips. 24 hours after seeding, samples were imaged directly using Zeiss Axio Observer Z1 or stained with NucBlue and fixed with 4% PFA for 10 minutes. Individual cell mCherry intensity was quantified on ImageJ using nuclear segmentation from 5 images per sample. Gaussian blur, thresholding, watershed, and analyze particle functions were applied to the nuclei and mCherry channel and to create individual selections for total cells and activated cells, respectively. The mCherry mask was applied to the corresponding mCherry image to measure the average fluorescence intensity within each activated cell. Percent activation was determined by the number of mCherry positive cells divided by the total number of nuclei. Activated mCherry intensity was calculated by averaging the mCherry intensity for cells above the defined threshold.

### Fibronectin-GFP Activation

For local activation, L929 anti-GFP synNotch receiver cells were seeded with 3T3 cells expressing FN-eGFP and nuclear localized miRFP703 in a ratio of 50:1 to a total of 5×10^4^ cells per well on a 8-well slide (Ibidi). After 3 days, cells were imaged on a Zeiss LSM780. For the titration experiments, cells were seeded in 8-well ibidi slides coated with 0.1% gelatin to a total of 3×10^4^ parental 3T3 cells and FN-GFP sender cells in following ratios 1:0, 50:1, 5:1, 2:1, 1:1 and 0:1 (Parental: FN-GFP). Following 8-10 days of culture, decellularization was performed following previously established methods (^43^). Briefly, cell-laden extracellular matrix was washed with PBS, wash buffers, and lysis buffer containing NP-40 for up to two hours to remove cellular debris. Removal of nuclear debris was confirmed by Hoechst staining prior to decellularization, which was used to monitor decellularization quality during the lysis phase of the protocol. Decellularized ECM was used immediately or stored at 4°C until use. 5-10×10^4^ L929 anti-GFP-synNotch receiver cells were seeded onto the decellularized matrices. After 2 days cells were imaged on a Zeiss LSM780 or Keyence BZ-X and on the same day analyzed by FACS on a Thermo Fisher Attune. For other co-culture experiments, L929 anti-GFP (or anti-mCherry) synNotch receiver cells were seeded at a 1:1 ratio with FN-GFP or FN-mCherry senders. Activation of engineered cells was imaged using Keyence BZ-X or fixed, stained for fibronectin (primary O/N 4°C, secondary 1 hour RT), and imaged on a Zeiss LSM780.

### Gelatin Hydrogel Surface Conjugation

Gelatin hydrogels were fabricated as previously described^33, 35^. Briefly, a 30W Epilog Mini 24 laser engraver (100% speed, 25% power, 2500 Hz) was used to cut a 150-mm polystyrene dish into 260-mm^2 hexagons. Each hexagon was masked with tape, and an inner circle was cut (18% speed, 6% power, 2500 Hz) and removed, exposing a polystyrene surface which was then treated with plasma (Harrick Plasma) for 10 minutes to improve gelatin adherence to polystyrene. Equal volumes of a 20% porcine gelatin solution (Sigma) and 8% MTG (Ajinomoto) solution were mixed and 200 ul were added to each coverslip. Flat or 10×10 um micromolded PDMS stamps were immediately applied to shape surface topography. After an overnight incubation to solidify, the hydrogels were rehydrated in water, and the stamp was removed. Coverslips were stored in PBS at 4°C until cell seeding.

PDMS stamps with 10×10 μm grooves of 2 μm height were fabricated with standard photolithography and soft lithography techniques^44^. Flat PDMS was used as a control substrate with no topography. A 1:1 ratio of GFP (500µg/ml, 200 µg/ml, and 20 µg/ml) and MTG (8% w/v) solution were added to a parafilm surface, and the gelatin coverslip was inverted onto the GFP- MTG droplet for 10 minutes^35^. The coverslips were then incubated for 1 hour at 37°C. Following incubation, the coverslips were washed 3 times with warm PBS.

L929 anti-GFP synNotch receiver cells were seeded at a density of 350,000 cells per coverslip and cultured for 72 hours. To detach the cells from the gelatin for flow cytometry analysis, the gelatin hydrogels were minced with a sterile X-acto knife and incubated in a 4 mg/ml collagenase IV solution for 45 minutes at 37°C. Digested gelatin was then filtered with a 40 µm cell strainer.

C3H anti-GFPsynNotch MyoD expressing cells were seeded at a density of 500,000 cells per coverslip, and cultured for 7 days. Coverslips were washed three times with warm PBS, fixed with ice-cold methanol and immunostained with mouse alpha-actinin primary antibody (Sigma, A7811) at 1:200 dilution for two hours. Coverslips were then stained with the secondary antibody goat anti-mouse conjugated to Alexa Fluor 546 and 4′,6-diamidino-2-phenylindole (DAPI) at 1:200 dilutions for 2 hours. ProLong gold antifade mountant (Thermofisher) was used to mount cells on glass coverslips.

Myotube count was performed in ImageJ through size and intensity thresholding of the alpha-actinin signal. The total number of cells per image was calculated by size and intensity thresholding of the DAPI signal. Myogenic index was determined by dividing the number of co- localized nuclei within the alpha-actinin by the total number of nuclei in the field of view. Orientation of aligned and isotropic myotubes was determined using the OrientationJ distribution plugin.

### 3D Ligand Conjugation and Activation

GelMA Synthesis: Porcine Gelatin (175G Bloom, Sigma G2625) was dissolved at 10g in 100mL in 0.25M carbonate-bicarbonate buffer (pH 9) at 50°C under argon. 0.4mL methacrylic anhydride (Sigma 276685) was added dropwise with stirring (500 rpm) and reacted for 3 hours at 50°C. The reaction was then cooled to 40°C, adjusted with 6M HCl to pH 7.4, transferred to 12-14 kDa cutoff dialysis tubing (Fisher Scientific), dialyzed for 3 days at 40°C against 4L deionized water (changed twice daily), and lyophilized. GelMA was stored at -80°C until use. Degree of methacrylation was calculated using ^1^H-NMR compared to unmodified gelatin.

GelMA was dissolved at 1% w/v in PBS at 37°C. Methyltetrazine (mTz)-PEG5-NHS Ester (Click Chemistry Tools) was first dissolved at 8.8mM in DMSO and added to GelMA solution dropwise with stirring to 0.88mM final concentration. Mixture was reacted at 37°C overnight, transferred to 12-14 kDa cutoff dialysis tubing (Fisher Scientific), and dialyzed for 3 days at 40°C against 4L deionized water (changed twice daily), and lyophilized. Substitution was verified using ^1^H-NMR compared to unmodified GelMA and used to estimate percent substitution of methacrylate and methyltetrazine. Ligands (GFP and mCherry) were modified using trans-Cyclooctene (TCO)- PEG4-NHS Ester (Click Chemistry Tools) to generate GFP-TCO and mCherry-TCO. TCO-PEG4- NHS Ester was dissolved at 10mM in DMSO, added at a 20-molar excess to GFP and mCherry in PBS (1-3mg/mL), and reacted for 1 hour at room temperature in Eppendorf tubes with orbital rocking. Following, mixture was purified using Zeba Spin Desalting Columns (7k MWCO, ThermoFisher) following manufacturer instructions, sterile filtered, and aliquoted to 1mg/mL and stored at -80°C. Reactivity was verified by reacting GelMA-mTz at 1% w/v and GFP-TCO at 100µg/mL for one hour at 37°C, running on 4-20% SDS-PAGE gel (Bio-Rad and Coomassie blue staining (Invitrogen).

### Cell Encapsulation and Activation

GelMA-mTz at 1% w/v was reacted with GFP-TCO at 50-100µg/mL for one hour at 37°C. Following, 18% w/v GelMA solution and freshly prepared 25mg/mL Lithium phenyl-2,4,6- trimethylbenzoylphosphinate (LAP) was added to a final concentration of 5-10% w/v GelMA, 0- 100ug/mL GFP-TCO, and 0.25% LAP. Engineered L929 or C3H cell lines were resuspended in this solution at 5-10×10^6^ cells / mL. An array of 20µL droplets of cell-laden hydrogel solutions were pipetted between two 25x75mm glass slides that were thrice treated with GelSlick (Lonza) separated by 3D-printed 400μm insert. Gels were crosslinked for 90-180s at 25mW/cm^2^ using an Omnicure2000 and collimating adapter (Excelitas) and transferred to individual wells with DMEM complete. Following 30 minute incubation at 37°C, media was exchanged to fresh DMEM complete. Cell-laden hydrogels were cultured up to 14 days with media changes every 2 days. In the hydrogel patterning experiment, cell-laden hydrogel solutions containing 0 or 50μg/mL GFP were pipetted in proximity to each other to initiate limited contact immediately prior to crosslinking. This process led to a stable biphasic gel that was treated similarly as described above. Samples were imaged live with Zeiss Axio Observer Z1 or Zeiss LSM880 confocal microscope. Individual cell mCherry intensity was quantified on ImageJ using constitutive BFP-signal to segment individual cells. Gaussian blur, thresholding, watershed, and analyze particle functions were applied to the BFP channel to create individual selections for each cell. This mask was applied to the corresponding mCherry image to measure the average fluorescence intensity within each cell. Percent activation was determined by the number of BFP+ cells that had an mCherry intensity higher than a defined threshold. For spatial activation within 3D, line plots were quantified using the “Plot Profile” feature on inverted microscope images in ImageJ and normalized across the entire time course. Evaluation of cell viability was performed using Live/Dead Viability/Cytotoxicity Kit (ThermoFisher) following manufacturer’s instructions. Briefly, cell-laden gels were incubated in a 2 µM calcein AM and 4 µM EthD-1 solution in PBS for 45 minutes prior to imaging on a Zeiss LSM880.

In the co-culture encapsulation experiment, GFP-sender fibroblasts and anti-GFP synNotch receiver cells were co-encapsulated within 5% GelMA hydrogels at a fixed total cell concentration of 20×10^6^/mL. The ratio of each cell type was varied. Hydrogels were imaged on an LSM880 and the percent mCherry expression was evaluated using image analysis, where the BFP-signal was used to segment individual receiver cells.

Fibrinogen (Sigma) was dissolved at 5mg/mL in warmed PBS. Methyltetrazine (mTz)-PEG5-NHS Ester (Click Chemistry Tools) was first dissolved at 20mM in DMSO and added to the Fibrinogen solution dropwise to a 0.16mM final concentration. Solution was incubated at room temperature with manual rocking every 10 minutes for one hour, transferred to 12-14 kDa cutoff dialysis tubing (Fisher Scientific), and dialyzed overnight at 4°C against 12L deionized water, and lyophilized. Resulting Fibrinogen-mTz was stored at -80°C until use. For generation of Fibrinogen-mCherry and cell encapsulation, Fibrinogen-mTz was dissolved at 20mg/mL in PBS and incubated with 160µg/mL mCherry-TCO in PBS for one hour at 37°C. DMEM was added to bring to a final 10mg/mL Fibrinogen + 100 µg/mL mCherry and used to resuspend cell pellet at 5×10^6^ anti- mCherry synNotch receiver cells/mL. Immediately prior, 0.2U Thrombin / mg Fibrinogen was added to the solution and samples were allowed to gel for 10 minutes at 37°C. Following a 15 minute wash with DMEM to remove unbound mCherry, samples were transferred to the incubator for culture. Following 24 hours of culture, HSC NuclearMask Deep Red was added for one hour at 37°C to visualize nuclei before imaging on Zeiss Axio Observer Z1.

### Microcontact Printing

For all surfaces exept Fig. S6B (see below), PDMS-coated coverslips were UV-O treated for 3 minutes before being incubated in 10% APTES in ethanol for 2 hours at 50C. Coverslips were then washed with water before incubating in 2% glutaraldehyde solution in ethanol at room temperature for 1 hour^43^. Coverslips were then washed again and inverted onto a 150µL fibronectin droplet at 50 µg/mL concentration in distilled water. The coverslips were allowed to incubate on the fibronectin droplet in a parafilm sealed petri dish overnight at 4C. After the overnight FN incubation, GFP was microcontact printed over the FN coating using cylindrical isotropic stamps cut from a slab of PDMS using a biopsy punch (8 mm diameter). To determine the dose-dependent effect of GFP concentration, 0, 10, 50, 100, and 200 μg/mL GFP was coated onto isotropic stamps, which were then inverted onto fibronectin-coated coverslips. We then imaged and quantified the mCherry fluorescence on day 3. Using the optimal concentration of GFP (100 μg/mL), we also measured mCherry dynamics up to day 28.

For restricted adhesion (Fig. S6B), PDMS coated coverslips were UV-O treated for 8-minutes before being microcontact printed with a stamp coated with 100 µg/mL of GFP and 50 µg/mL of FN. After patterning, coverslips were incubated in 2% pluronic in distilled water for 15 minutes at room temperature before being washed with PBS and seeded with cells.

To generate micropatterns of GFP, Solidworks was used to design desired patterns (square arrays ranging from 100 μm to 1 mm, concentric circles, aligned rows, and letters), which were then printed using a digital light processing (DLP) 3-D printer (CADworks3D). After 3D printing, the replica molds were placed in 200 proof IPA overnight to ensure all uncured resin was removed. The molds were then UV cured for 1 hour (back side for 20 minutes, feature side for 40 minutes) to finalize the curing process. PDMS (Sylgard 187) was poured into the molds and placed in a vacuum chamber for 30 minutes to degas bubbles. The molds were then placed in a 65°C oven to cure the PDMS overnight. After curing, PDMS stamps were removed from the molds in preparation for microcontact printing.

Microcontact printing with PDMS stamps molded on 3-D printed templates was performed the same as with isotropic stamps. Stamps were incubated with 200 μg/mL GFP for 24h.

For perpendicular row patterns, one stamp was coated with GFP (200 μg/mL) and another stamp was coated with mCherry (200 μg/mL). These stamps were manually positioned sequentially in a perpendicular orientation. In this experiment, a monoclonal population of dual receiver fibroblasts was used. Patterns were stored dry at 4°C until use, and incubated in DMEM+10% FBS for a minimum of 1 hour prior to cell seeding. Cells were seeded on patterns at 5-20×10^4 cells/cm^2.

### Dual Ligand Patterning with Capillary Microfluidic Device

Solidworks was used to design a 4-row capillary fluidic device with two disconnected inlets. Shallow channels (100µm distance from the substrate) were designed to guide protein solutions. These shallow channels were surrounded by 1 mm deep channels, intended as voids. The inverse design was 3-D printed using a DLP printer (CADWorks), which was then replica molded in PDMS. Inlets were created using 1.5mm biopsy punches and air ventilation punches were created on two opposite sides ends of the device to allow optimal pressure for capillary fluid transfer. Prior to protein patterning, the feature side surface of the device was UV plasma treated for 7 seconds, creating a hydrophilic surface for capillary action. The device was then placed facedown onto tissue culture treated Ibidi 2-wells. GFP (500µg/mL) and mCherry (1000µg/mL) were then pipetted into separate inlets and filled their respective shallow channels. The device was then placed into a petri dish and parafilm sealed before overnight incubation at 4°C. The next day, the device was incubated without parafilm at room temperature for 15 minutes. The fluidic device was then carefully removed from the Ibidi wells to minimize liquid disruption. The Ibidi well was left at room temperature for 15 minutes for the protein solutions to dry. The Ibidi well was then UV treated under the biosafety cabinet for 1 hour to sterilize. DMEM with 10% FBS was pipetted into the wells and incubated for 1 hour before cell seeding. Dual-lineage cells were seeded at 1.9×10^5^/cm^2^ and cultured for three days prior to fixation and staining for α-actinin and VEGFR2.

### Data Quantification

We studied the spatial control dynamics over time by measuring the Pearson’s Coefficient on days 2, 5, and 10. This was done using the JACoP plugin in ImageJ by comparing the binary mask to the mCherry channel. Line plots were quantified using the “Plot Profile” feature in ImageJ and normalized to individual images. The number of nuclei in the field of view of each image were counted, and the number of myotubes was counted using ImageJ. Myogenic index was calculated through dividing the number of nuclei within each myotube by the total number of nuclei. The GFP channel was used to determine which nuclei were on and off-pattern. For calculating on-pattern myogenic index, all nuclei outside the GFP pattern were excluded. The sarcomeric α-actinin mask was overlaid on the on-pattern nuclei and used to calculate the myogenic index. Similarly, to quantify off-pattern myogenic index, we excluded all nuclei located within the GFP patterns and used the same sarcomeric α-actinin mask to measure the myogenic index. For Coherency quantification, 200 and 500µm rows and curves were thresholded using the same methods used to create the myotube mask, except instead of using the thresholded image to create a selection/mask, we quantified the thresholded myotube image itself. The OrientationJ plugin on ImageJ was used to quantify the coherency of all patterns. Since curved rows are not straight, we need to straighten them to get a fair quantification of how the myotubes align with the curves. A fragmented line was drawn manually following the GFP pattern of the curve and used to straighten the myotube threshold image before quantifying the coherency. All data was processed in GraphPad Prism 9 and validated with statistics.

### Plate-drying of Ligand

For single-ligand activation, ligands (mCherry and GFP) were prepared at 100μg/mL in sterile DI water and added at 15μg/cm^2^. For dual-ligand patterning, 8uL droplets of ligand at 200μg/mL in sterile DI water were deposited in distinct regions within each well. Plates were left to dry in the biosafety cabinet overnight, protected from light and then washed once with PBS prior to cell seeding. Anti-mCherry synNotch ETV2-BFP or dual-lineage fibroblasts were seeded at 5- 20×10^4^ cells/cm^2^ and cultured for three days prior to flow cytometry analysis or fixation and staining for VEGFR2.

### Staining

Flow Cytometry: cells were detached using TrypLE (ThermoFisher) and washed once prior to incubation with fluorescently-tagged antibodies in PBS+5%FBS for 30 minutes-1 hour at 4°C. Following, cells were washed twice with PBS+5%FBS and filtered through 35μm cell strainer prior to analysis with ARIA II.

Following culture, cells were washed once with PBS, fixed with 4% paraformaldehyde or 10% ice-cold methanol for 10 minutes and then washed 3x with PBS for 5 minutes each. Samples were stained immediately or further permeabilized with 0.1% Triton X-100 in PBS for 5-10 minutes and then washed 3x with PBS for 5 minutes. Cells were blocked for 1 hour with 2% BSA at room temperature, then incubated with primary antibodies for 2 hours at room temperature or overnight at 4°C. Following three washes with PBS, samples were incubated with secondary antibodies for one hour at room temperature, then washed again prior to imaging directly or staining nuclei with NucBlue (15 minutes, ThermoFisher) or Nuclear Mask Deep Red (30 min, ThermoFisher). Samples on coverslip were mounted with gold-antifade mounting solution (ThermoFisher).

### Imaging/Microscopy

Unless otherwise stated, a digital Microscope (Keyence BZ-X) was used to image experiments. Tiling was done with the built-in Keyence software. All images within individual experiments were taken with the same settings (Light strength, exposure, No LUT). BFP, GFP, and mCherry signals were captured using the respective Filter cubes: BFP, GFP, TexasRed, Cy5-NX.

### Statistics

Individual data points in graphs represent distinct samples. Statistics were calculated in Prism, using Unpaired T-test two-tailed or one-way Anova between groups. * p<.05, ** p=<.01, *** p=<.001, **** p=<.0001

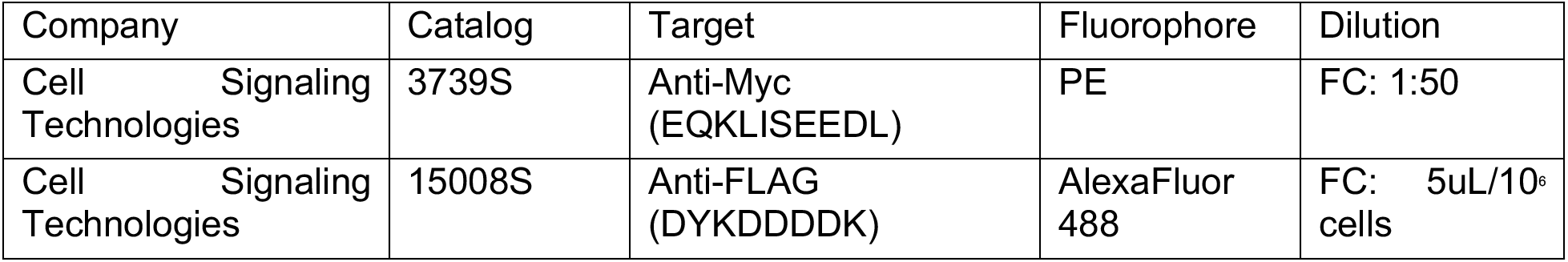
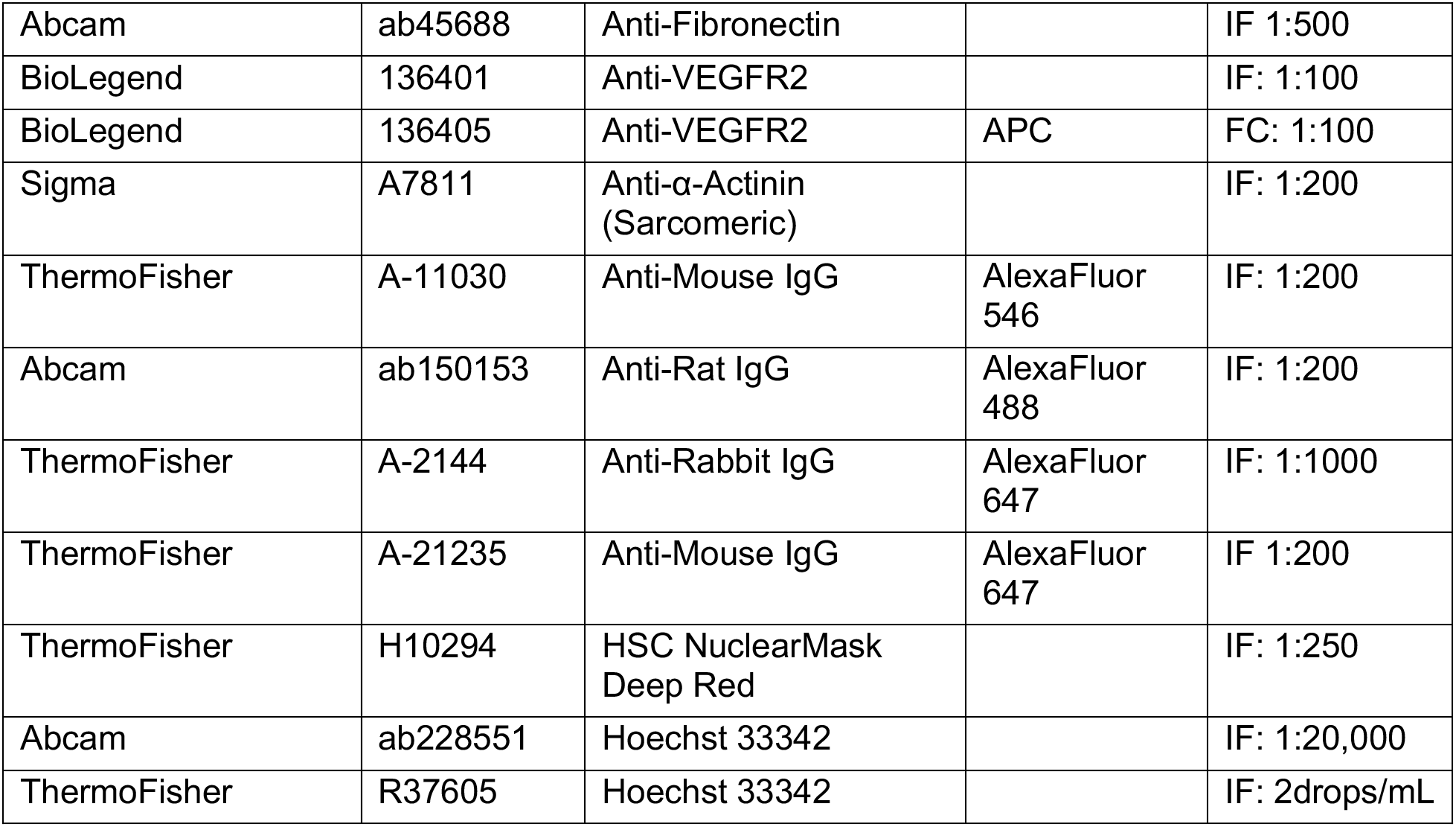

| Company | Catalog | Target | Fluorophore | Dilution |
| --- | --- | --- | --- | --- |
| Cell Signaling Technologies | 3739S | Anti-Myc (EQKLISEEDL) | PE | FC: 1:50 |
| Cell Signaling Technologies | 15008S | Anti-FLAG (DYKDDDDK) | AlexaFluor 488 | FC: 5uL/10<sup>6</sup> cells |
| Abcam | ab45688 | Anti-Fibronectin |  | IF 1:500 |
| BioLegend | 136401 | Anti-VEGFR2 |  | IF: 1:100 |
| BioLegend | 136405 | Anti-VEGFR2 | APC | FC: 1:100 |
| Sigma | A7811 | Anti-α-Actinin<br>(Sarcomeric) | | IF: 1:200 |
| ThermoFisher | A-11030 | Anti-Mouse IgG | AlexaFluor<br>546 | IF: 1:200 |
| Abcam | ab150153 | Anti-Rat IgG | AlexaFluor<br>488 | IF: 1:200 |
| ThermoFisher | A-2144 | Anti-Rabbit IgG | AlexaFluor<br>647 | IF: 1:1000 |
| ThermoFisher | A-21235 | Anti-Mouse IgG | AlexaFluor<br>647 | IF 1:200 |
| ThermoFisher | H10294 | HSC NuclearMask<br>Deep Red |  | IF: 1:250 |
| Abcam | ab228551 | Hoechst 33342 |  | IF: 1:20,000 |
| ThermoFisher | R37605 | Hoechst 33342 |  | IF: 2drops/mL |

