## Supplemental Figures and Figure legends for "Engineering Programmable Material-To-Cell Pathways Via Synthetic Notch Receptors To Spatially Control Cellular Phenotypes In Multi-Cellular Constructs"

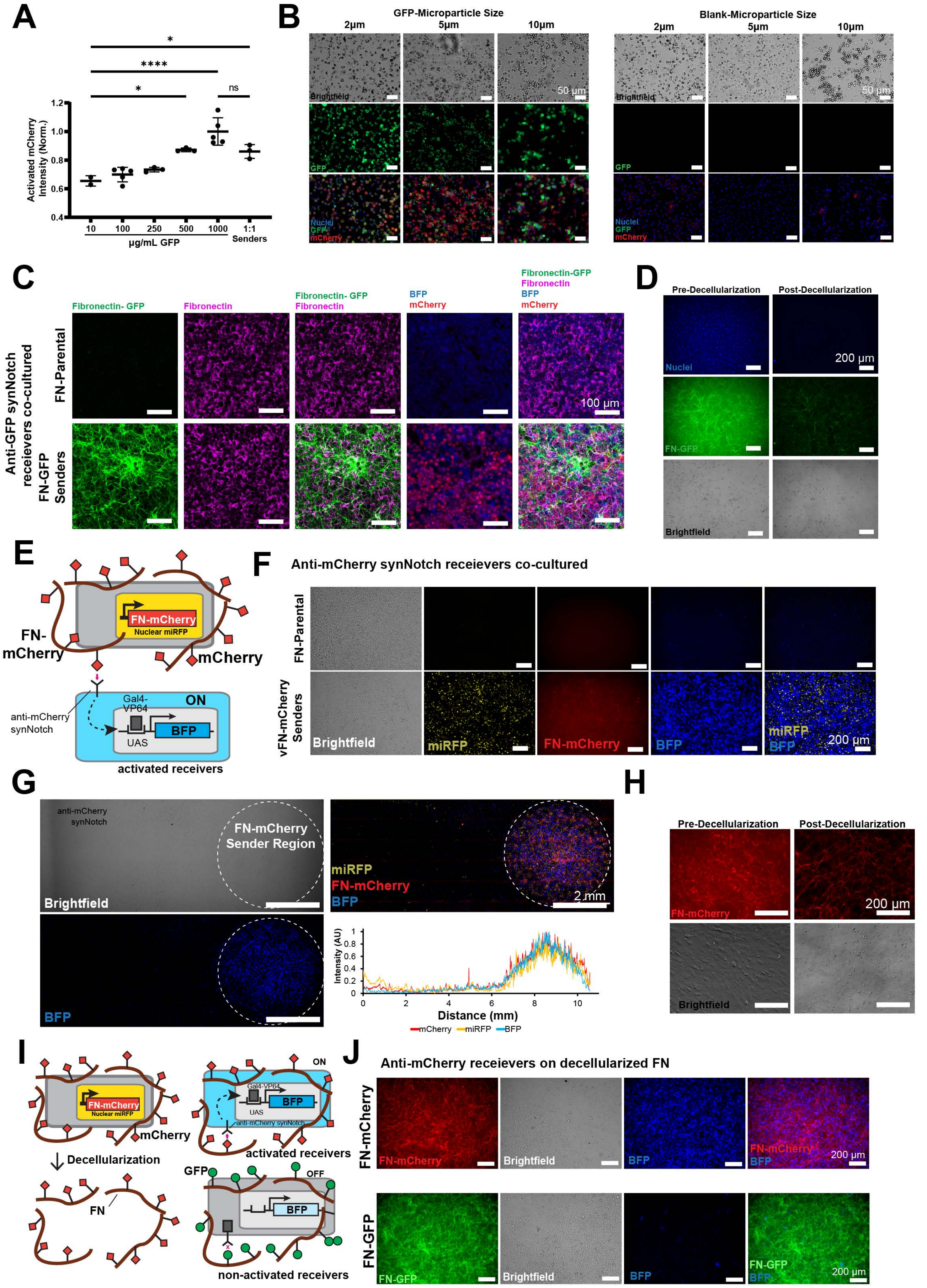

A
Activated mCherry Intensity (Norm.)
*
****
*
ns
10
100
250
500
1000
1:1 Senders
µg/mL GFP
B
GFP-Microparticle Size
Blank-Microparticle Size
2µm
5µm
10µm
Brightfield
GFP
Nuclei
mCherry
50 µm
C
Anti-GFP synNotch receievers co-cultured
FN-Parental
FN-GFP Senders
Fibronectin- GFP
Fibronectin
BFP
mCherry
100 µm
D
Pre-Decellularization
Post-Decellularization
Nuclei
FN-GFP
Brightfield
200 µm
E
FN-mCherry
Nuclear miRFP
mCherry
anti-mCherry synNotch
Gal4-VP64
UAS
BFP
ON
activated receivers
F
Anti-mCherry synNotch receievers co-cultured
FN-Parental
vFN-mCherry Senders
Brightfield
miRFP
FN-mCherry
BFP
200 µm
G
anti-mCherry synNotch
FN-mCherry Sender Region
Brightfield
miRFP
FN-mCherry
BFP
2 mm
Intensity (AU)
Distance (mm)
mCherry
miRFP
BFP
H
Pre-Decellularization
Post-Decellularization
FN-mCherry
Brightfield
200 µm
I
FN-mCherry
Nuclear miRFP
mCherry
Decellularization
FN
ON
Gal4-VP64
BFP
UAS
anti-mCherry synNotch
activated receivers
GFP
OFF
non-activated receivers
J
Anti-mCherry receievers on decellularized FN
FN-mCherry
FN-GFP
Brightfield
BFP
200 µm

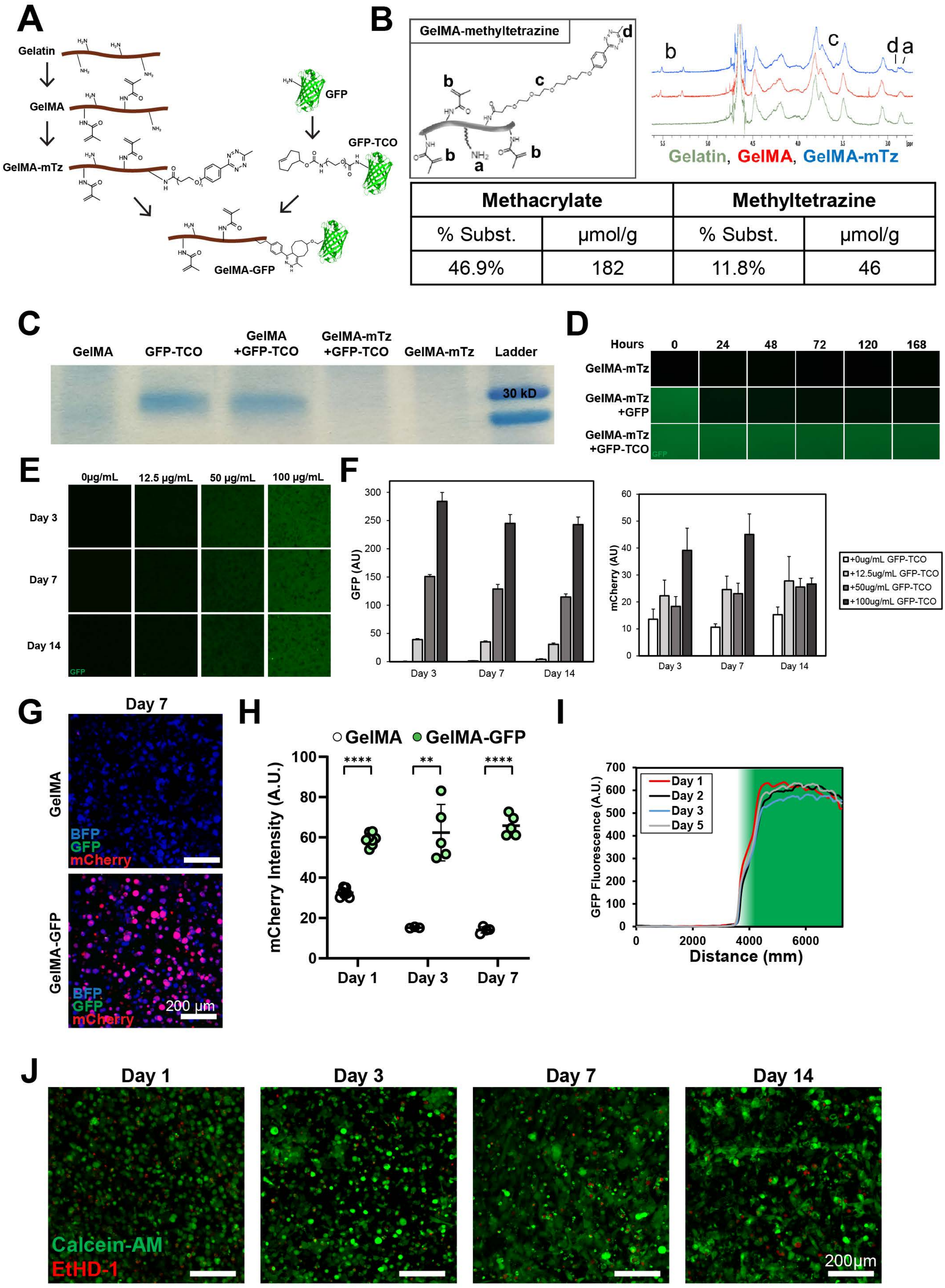


| Methacrylate | | Methyltetrazine | |
|---|---|---|---|
| % Subst. | μmol/g | % Subst. | μmol/g |
| 46.9% | 182 | 11.8% | 46 |

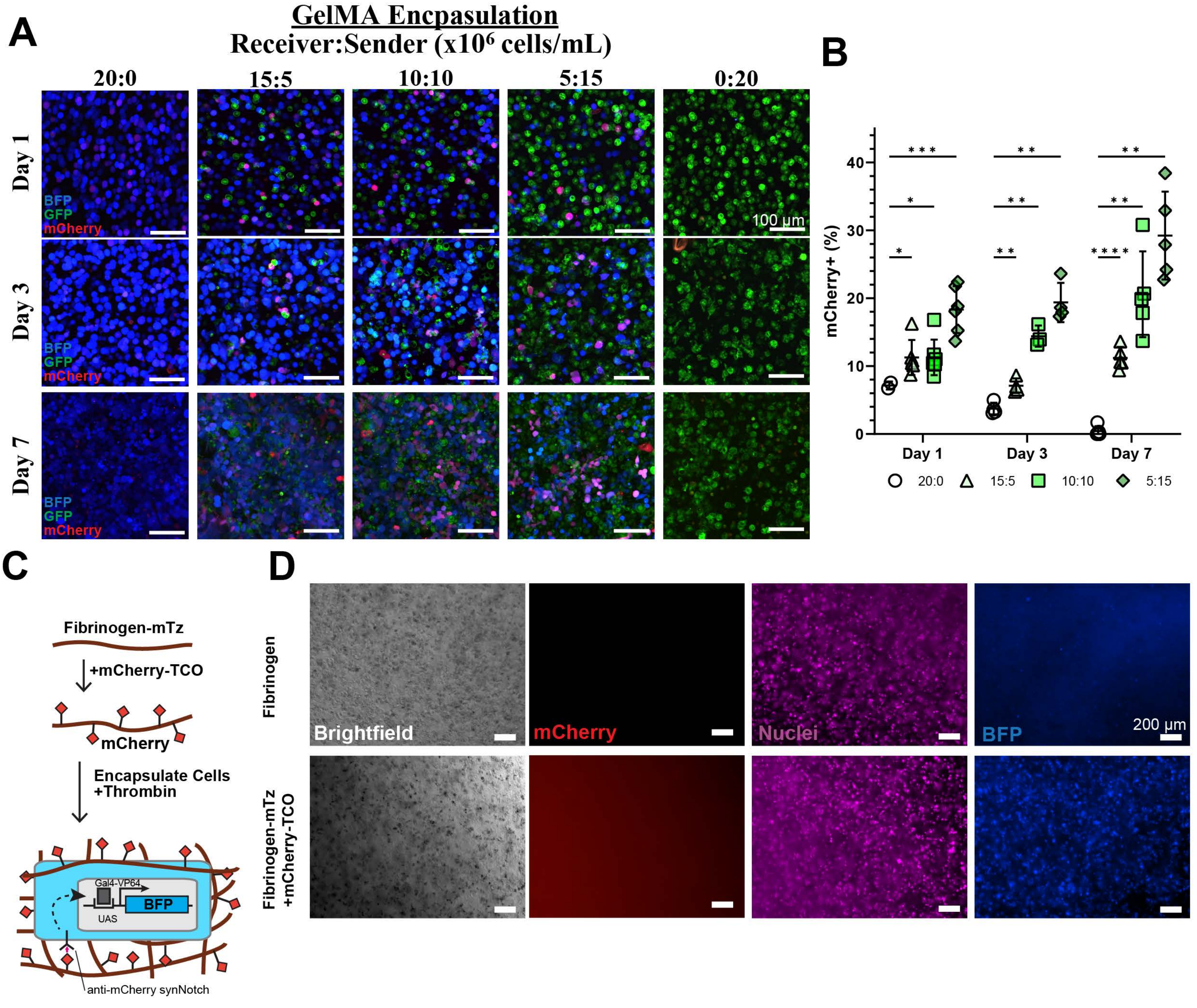
A
GelMA Encpasulation
Receiver:Sender (x10⁶ cells/mL)
20:0
15:5
10:10
5:15
0:20
Day 1
Day 3
Day 7
BFP
GFP
mCherry
100 µm
B
mCherry+ (%)
Day 1
Day 3
Day 7
20:0
15:5
10:10
5:15
C
Fibrinogen-mTz
+mCherry-TCO
mCherry
Encapsulate Cells
+Thrombin
Gal4-VP64
UAS
BFP
anti-mCherry synNotch
D
Fibrinogen
Fibrinogen-mTz
+mCherry-TCO
Brightfield
mCherry
Nuclei
BFP
200 µm

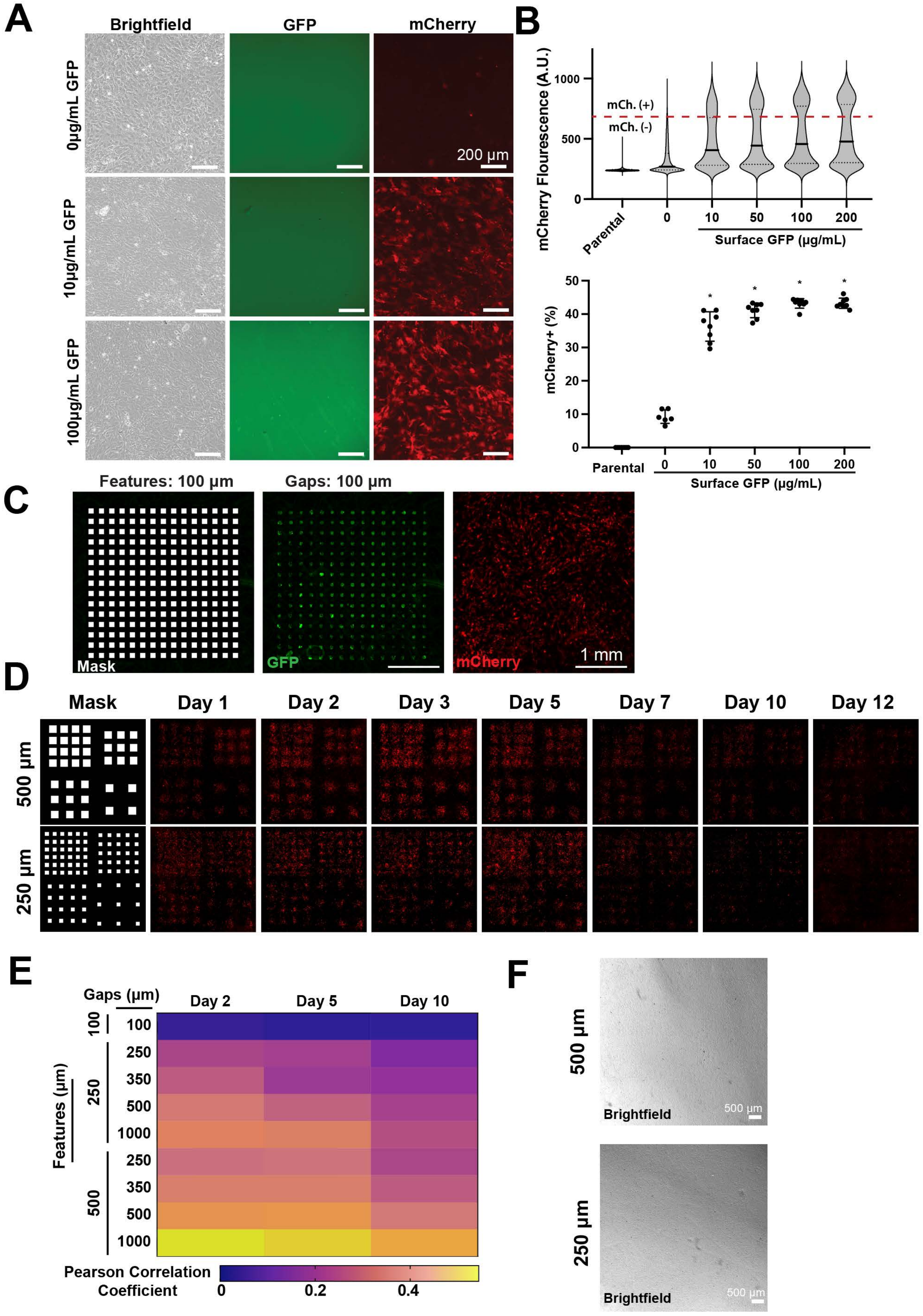

A
Brightfield
GFP
mCherry
0μg/mL GFP
10μg/mL GFP
100μg/mL GFP
200 μm
B
mCherry Flourescence (A.U.)
mCh. (+)
mCh. (-)
Parental
Surface GFP (μg/mL)
mCherry+ (%)
C
Features: 100 μm
Gaps: 100 μm
Mask
GFP
mCherry
1 mm
D
Mask
Day 1
Day 2
Day 3
Day 5
Day 7
Day 10
Day 12
500 μm
250 μm
E
Gaps (μm)
Day 2
Day 5
Day 10
Features (μm)
Pearson Correlation Coefficient
F
Brightfield
500 μm
250 μm

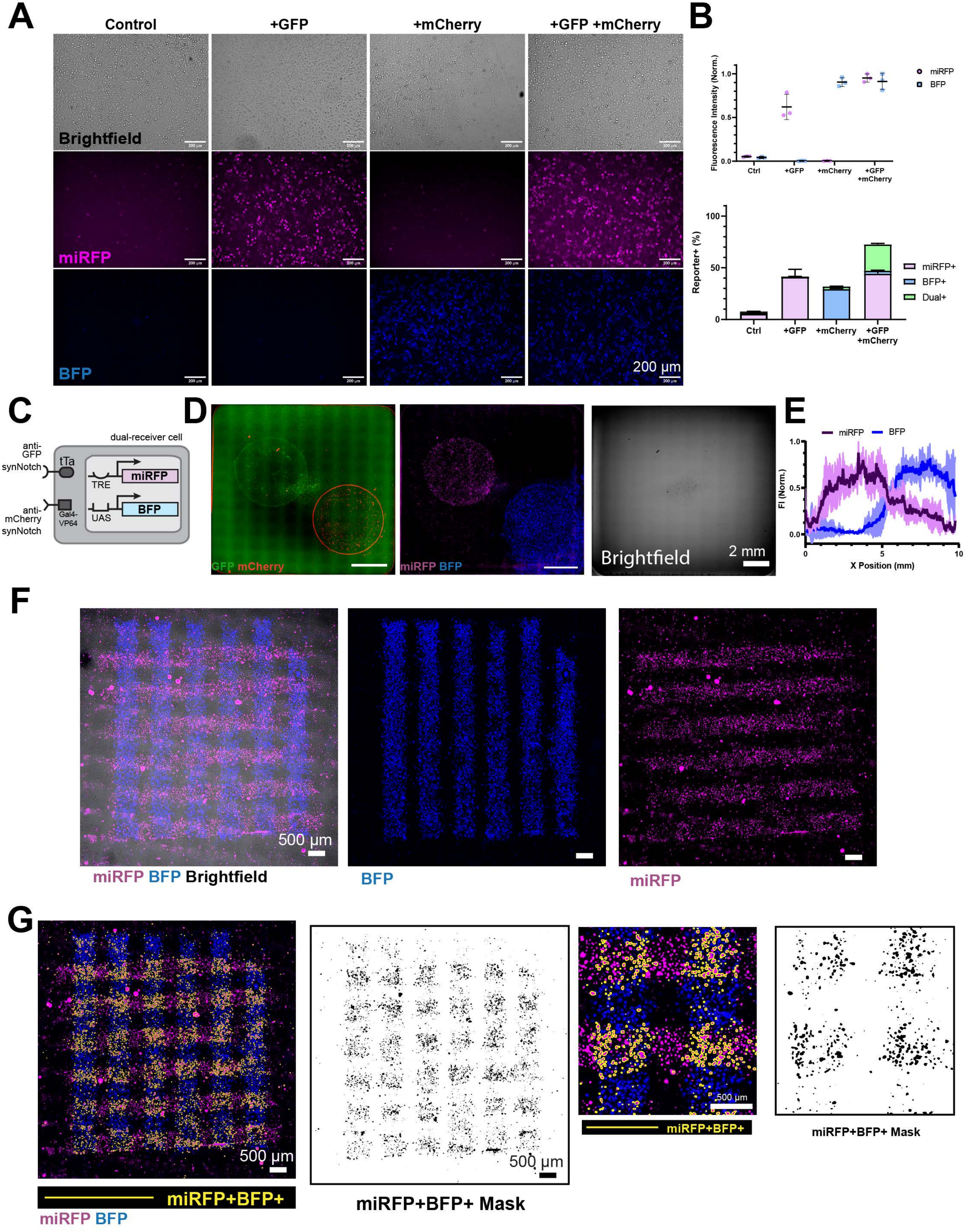

A
Control
+GFP
+mCherry
+GFP +mCherry
Brightfield
miRFP
BFP
200 µm
B
Fluorescence Intensity (Norm.)
miRFP
BFP
Ctrl
+GFP
+mCherry
+GFP +mCherry
Reporter+ (%)
miRFP+
BFP+
Dual+
C
dual-receiver cell
anti-GFP synNotch
tTa
TRE
miRFP
anti-mCherry synNotch
Gal4-VP64
UAS
BFP
D
GFP mCherry
miRFP BFP
Brightfield
2 mm
E
miRFP
BFP
FI (Norm.)
X Position (mm)
F
500 µm
miRFP BFP Brightfield
BFP
miRFP
G
500 µm
miRFP+BFP+
miRFP BFP
miRFP+BFP+ Mask

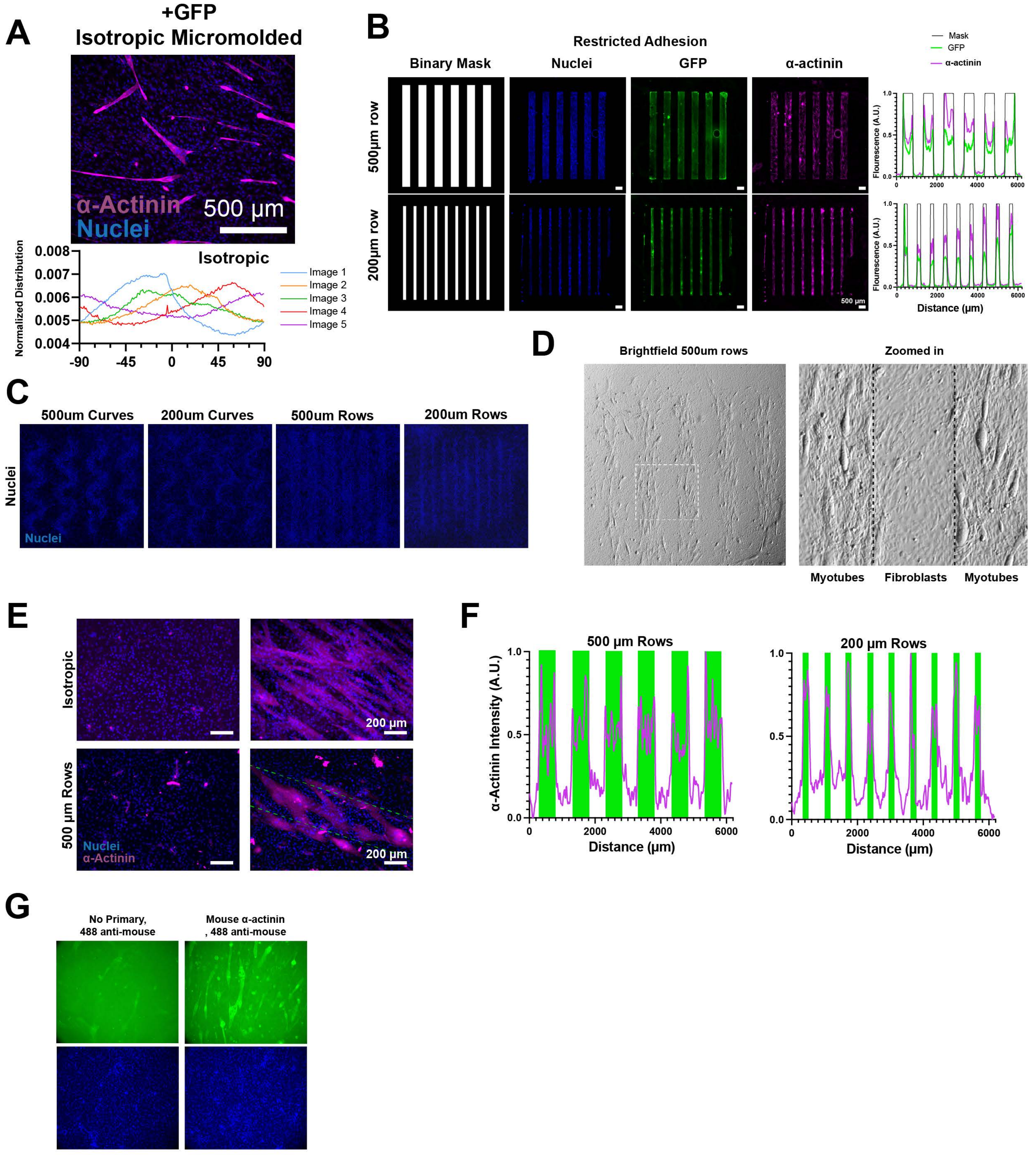

A
+GFP
Isotropic Micromolded
α-Actinin
Nuclei
500 μm
Normalized Distribution
Isotropic
Image 1
Image 2
Image 3
Image 4
Image 5
B
Restricted Adhesion
Binary Mask
Nuclei
GFP
α-actinin
500μm row
200μm row
500 μm
Mask
GFP
α-actinin
Flourescence (A.U.)
Distance (μm)
C
500um Curves
200um Curves
500um Rows
200um Rows
Nuclei
D
Brightfield 500um rows
Zoomed in
Myotubes
Fibroblasts
Myotubes
E
Isotropic
500 μm Rows
Nuclei
α-Actinin
200 μm
F
500 μm Rows
200 μm Rows
α-Actinin Intensity (A.U.)
Distance (μm)
G
No Primary,
488 anti-mouse
Mouse α-actinin
, 488 anti-mouse

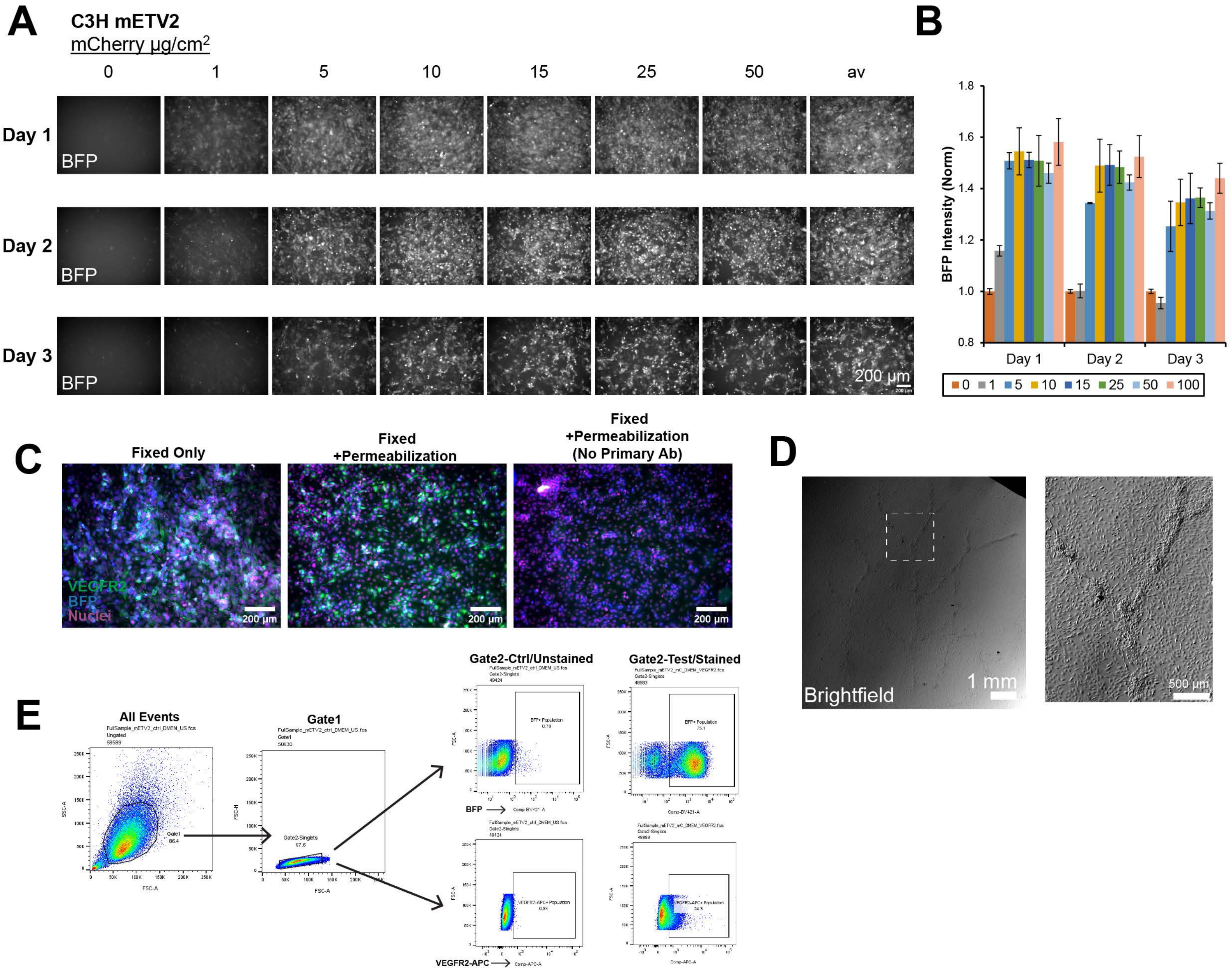
A
C3H mETV2
mCherry µg/cm2
0
1
5
10
15
25
50
av
Day 1
Day 2
Day 3
BFP
200 µm
B
BFP Intensity (Norm)
1.8
1.6
1.4
1.2
1.0
0.8
Day 1
Day 2
Day 3
0 1 5 10 15 25 50 100
C
Fixed Only
Fixed +Permeabilization
Fixed +Permeabilization (No Primary Ab)
VEGFR2
BFP
Nuclei
200 µm
D
Brightfield
1 mm
500 µm
E
All Events
Gate1
Gate2-Ctrl/Unstained
Gate2-Test/Stained
BFP+ Population
VEGFR2-APC+ Population
BFP
VEGFR2-APC
SSC-A
FSC-A
FSC-H
Gate2-Singlets
97.6
86.4

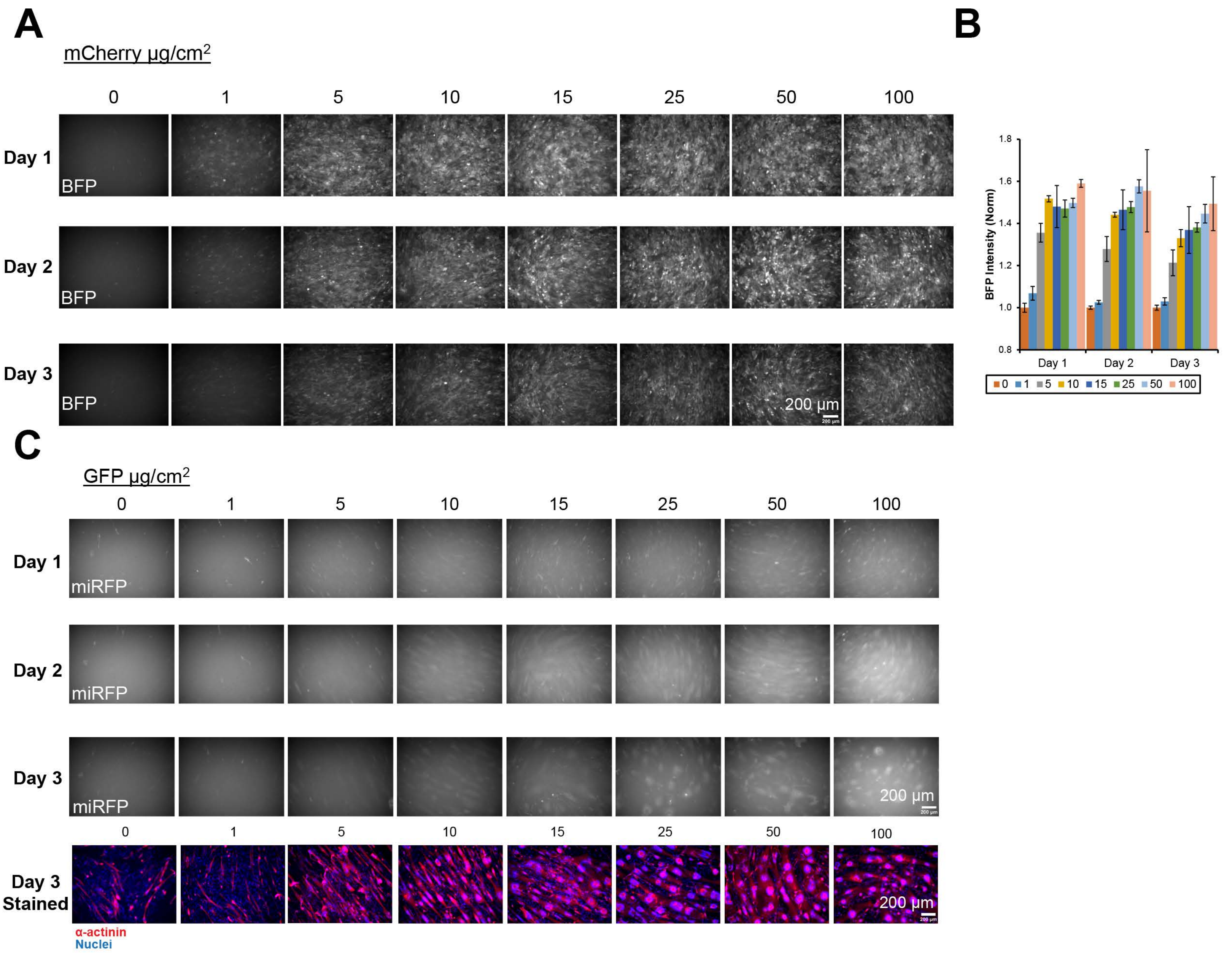
A
mCherry μg/cm2
0
1
5
10
15
25
50
100
Day 1
Day 2
Day 3
BFP
200 μm
B
BFP Intensity (Norm)
1.8
1.6
1.4
1.2
1.0
0.8
Day 1
Day 2
Day 3
0
1
5
10
15
25
50
100
C
GFP μg/cm2
0
1
5
10
15
25
50
100
Day 1
Day 2
Day 3
miRFP
200 μm
Day 3 Stained
α-actinin
Nuclei

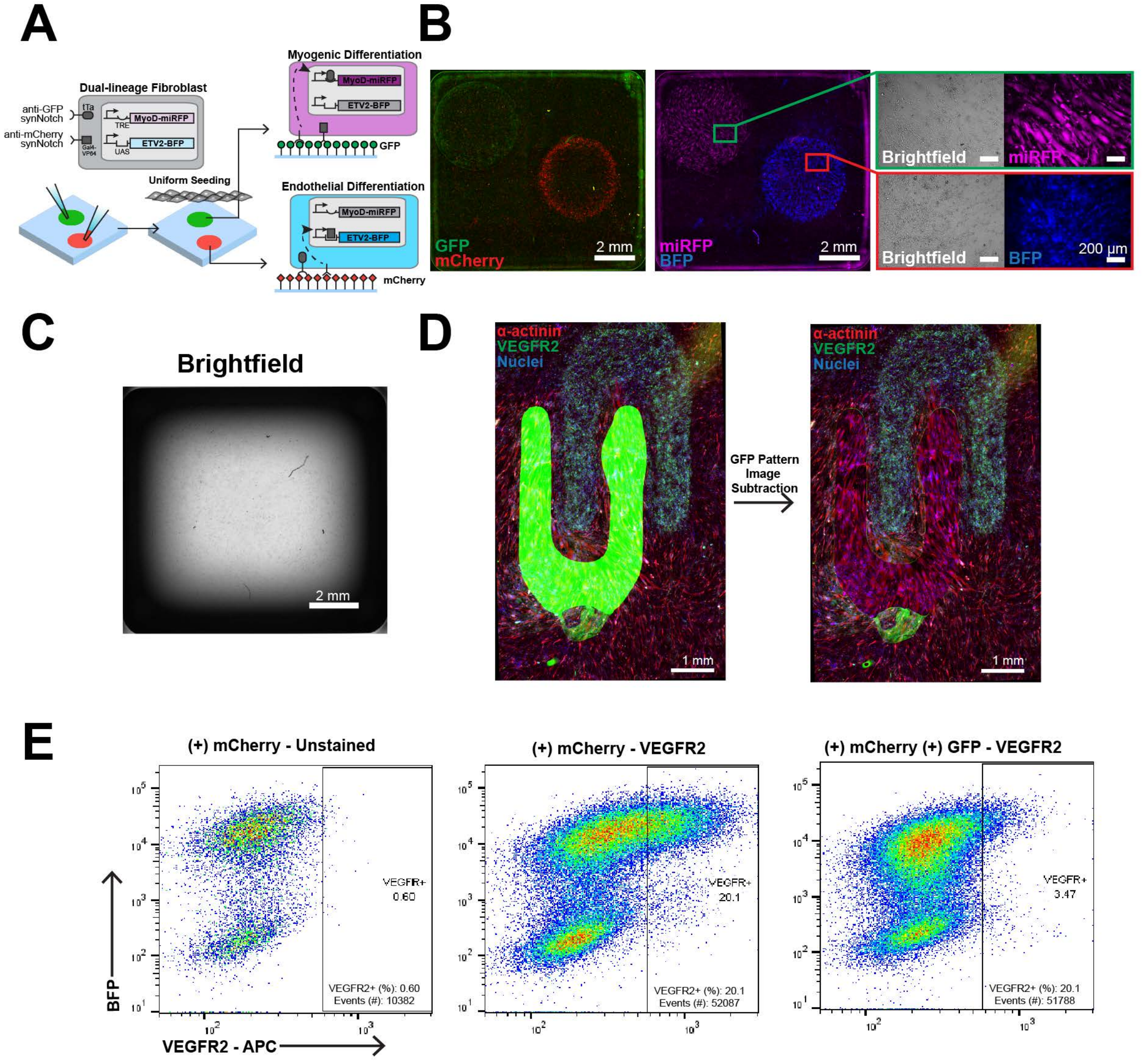
A
Dual-lineage Fibroblast
anti-GFP synNotch
anti-mCherry synNotch
tTa
TRE
MyoD-miRFP
Gal4-VP64
UAS
ETV2-BFP
Uniform Seeding
Myogenic Differentiation
MyoD-miRFP
ETV2-BFP
GFP
Endothelial Differentiation
MyoD-miRFP
ETV2-BFP
mCherry
B
GFP
mCherry
2 mm
miRFP
BFP
2 mm
Brightfield
miRFP
Brightfield
BFP
200 µm
C
Brightfield
2 mm
D
α-actinin
VEGFR2
Nuclei
1 mm
GFP Pattern Image Subtraction
α-actinin
VEGFR2
Nuclei
1 mm
E
(+) mCherry - Unstained
VEGFR+
0.60
VEGFR2+ (%): 0.60
Events (#): 10382
(+) mCherry - VEGFR2
VEGFR+
20.1
VEGFR2+ (%): 20.1
Events (#): 52087
(+) mCherry (+) GFP - VEGFR2
VEGFR+
3.47
VEGFR2+ (%): 20.1
Events (#): 51788
BFP
VEGFR2 - APC

### SUPPLEMENTAL FIGURE LEGENDS

**Figure S1.** (A) Average mCherry intensity of activated reporter cells quantified by image analysis following 24-hour co-culture with 5μm GFP microparticles of different conjugation concentrations or GFP sender cells. Data represents mean ± s.d, n=3-5, p>.05 (ns), p<0.05 (*), p<0.0001(****). (B) Fluorescence and brightfield images of GFP-conjugated (left) and Blank (right) microparticles of varying sizes following 24-hour co-culture with engineered receiver fibroblasts. Scale bars, 50 μm. (C) Fluorescence images of anti-GFP synNotch receiver fibroblasts with 3T3 parental fibroblasts (top) or Fibronectin-GFP (FN-GFP) sender fibroblasts co-cultured for 72 hours prior to fibronectin immunostaining with anti-fibronectin and Alexa-647 antibodies. Scale bars, 100 μm. (D) Comparison of the presence of nuclei (blue) and FN-GFP (green) before and after the decellularization process. Scale bars, 200 μm. (E) Schematic of Fibronectin-mCherry (FN-mCherry) producing sender fibroblasts, with a miRFP nuclear tag co-cultured with anti-mCherry synNotch/Gal4 receiver fibroblasts that induce the expression of BFP reporter. (F) Fluorescence images of anti-mCherry synNotch receiver fibroblasts co-cultured with parental fibroblasts (top) or FN-mCherry sender fibroblasts (bottom) for 48 hours. Scale bars, 200 μm. (G) Brightfield and fluorescence images of anti-mCherry synNotch reporter fibroblasts uniformly seeded onto a local region of FN-mCherry sender fibroblasts, which were seeded 30 minutes prior within a small droplet (white line indicates region of FN-mCherry seeding). Line plot represents the normalized fluorescence intensity across the x-axis. Scale bars, 2mm. (H) Fluorescence and brightfield images comparing FN-mCherry before and after the decellularization process. Scale bars, 200μm. (I) Schematic detailing FN-mCherry sender culture, followed by decellularization, then reseeding with anti-mCherry receivers. (J) Fluorescence images of anti-mCherry synNotch reporter fibroblasts cultured on decellularized FN-mCherry ECM (top) or FN-GFP ECM (bottom) taken 48 hours following seeding.

**Figure S2.** (A) Schematic of covalent substitutions to Gelatin, with methacrylate and methyltetrazine, and GFP, with trans-Cyclooctene, to generate gelatin methacryloyl (GelMA) conjugated to GFP (GelMA-GFP) (B) Functional groups of GelMA-methyltetrazine (GelMA-mTz) with corresponding NMR peaks during each step of gelatin modification. Percent substitution and molar content of methacrylate group and methyltetrazine estimated with $^{1}$H-NMR. (C) Coomassie-blue stained SDS-PAGE gel of varying combinations of GelMA±mTz with GFP±trans-Cyclooctene (TCO) to determine conjugation. GelMA±mTz is faintly visible due to polydisperse molecular weight. Darker colored band represents GFP (MW: 27 kDa). Protein ladder (right) shown for 30 kDa reference. (D) Fluorescence images of GFP signal within GelMA-mTz hydrogels loaded with either no GFP, GFP, or GFP-TCO incubated in PBS at 37°C for up to 168 hours. (E) Fluorescence images of GFP signal of anti-GFP synNotch receiver fibroblast-laden GelMA-mTz hydrogels loaded with varying concentrations of GFP-TCO (0, 12.5, 50, 100μg/mL) prior to photocrosslinking. (F) Image analysis based quantifications of GFP intensity and activated anti-GFP synNotch receiver fibroblasts (mCherry signal) within GelMA-mTz hydrogels containing 0, 12.5, 50, or 100μg/mL GFP taken up to 14 days. Data represents mean ± s.d, n=3 (G) Z-projected fluorescence images of anti-GFP synNotch mCherry fibroblasts encapsulated within GelMA-mTz hydrogels containing 0 or 50ug/mL GFP-TCO at Day 7. Scale bars, 200μm. (H) mCherry Intensity of encapsulated anti-GFP synNotch mCherry fibroblasts quantified by image analysis following 1, 3, and 7 days of culture within GelMA or GelMA-GFP hydrogels. Data represents mean ± s.d, n=3-8, p<0.01 (**), p<0.0001(****). (I) Plot profile of normalized GFP intensity distribution across the length of the bi-phasic GelMA hydrogel, where one portion contains GFP, 1, 2, 3, and 5 days after encapsulation. Green bar indicates the region containing GFP. (J) Fluorescence images of anti-GFP synNotch fibroblast-laden GelMA hydrogels stained with Live/Dead taken 1, 3, 7, and 14 days after fabrication. Live cells are stained with Calcein-AM, dead cells are stained with ethidium homodimer-1 (EtHD-1). Scale bars, 200μm.

**Figure S3.** (A) Z-projected fluorescence images of anti-GFP synNotch fibroblasts with inducible mCherry co-encapsulated with GFP sender fibroblasts in varying ratios (20:0, 15:5, 10:10, 5:15, 0:20 Receivers:Senders), taken 1, 3, and 7 days following encapsulation. Scale bars, 100μm. (B) Percent of mCherry expressing receiver cells quantified by image analysis following 1, 3, and 7 days of culture with varying ratios of sender cells. Data represents mean ± s.d, n=3-5, $p<.05$(*), $p<.01$(**), $p<.001$(***), $p<0.0001$(****). (C) Schematic of Fibrinogen-mTz and mCherry-TCO reaction to generate Fibrinogen-mCherry used to encapsulate anti-mCherry reporter fibroblasts with inducible BFP. (D) Fluorescence images of anti-mCherry reporter fibroblasts encapsulated within Fibrinogen or Fibrinogen-mTz with mCherry-TCO for 24 hours. Nuclei are stained with HSC NuclearMask Deep Red. Scale bars, 200μm.

**Figure S4.** (A) Day 2 fluorescence images showing mCherry activation in anti-GFP reporter fibroblasts on no GFP, 10μg/mL microcontact printed GFP, and 100μg/mL microcontact printed GFP. Scale bars, 200μm. (B) Violin plot of mCherry intensity, quantified with flow cytometry, dose response to 0, 10, 50, 100, and 200μg/mL concentrations of microcontact printed GFP after 48 hours. Dotted line indicates the threshold value to designate mCherry-positive cell. Percent of mCherry expressing cells quantified by flow cytometry 48 hours after seeding onto GFP microcontact printed substrate with varying GFP concentrations. Data represents mean ± s.d, n=6-8, $p<0.05$(*) (C) Microcontact printed 100μm GFP squares with 100μm interspace length and resulting mCherry activation signals taken two days after seeding. Scale bars, 1mm. (D) Fluorescence images of mCherry activation from days 1-12 on 250 and 500μm width GFP squares with varying interspace lengths. (E) Heatmap demonstrating Pearson correlation coefficient between mCherry and GFP signals of each square width and interspace length at Days 2, 5, and 10. Color map represents the mean coefficient, n=7-8. (F) Brightfield images of anti-GFP reporter fibroblasts 2 days following uniform seeding onto 500 and 250μm GFP squares. Scale bars, 500μm.

**Figure S5.** (A) Fluorescence images of anti-GFP and anti-mCherry dual-reporter fibroblasts in the presence of no ligand (control) or plate-dried GFP, mCherry, or both GFP and mCherry. Images taken 24 hours following seeding. Scale bars, 200μm. (B) Quantified fluorescence intensity of miRFP and BFP reporters and percent of cells expressing each or both reporters, quantified via flow cytometry, in the presence or absence of passively adsorbed GFP and/or mCherry. Data represents mean ± s.d, n=3. (C) Schematic of dual-reporter L929 cell where anti-GFP synNotch drives miRFP reporter gene and anti-mCherry synNotch orthogonally activates BFP (D) Fluorescence images of plate-dried GFP and mCherry droplets and subsequent dual reporter expression and brightfield taken 48 hours after uniform seeding of engineered fibroblasts. Scale bars, 2mm. (E) Normalized plot profiles of miRFP and BFP intensity across the x-axis 48 hours following seeding onto GFP and mCherry droplet pattern. Line profiles represent mean ± s.d, n=4. (F) Fluorescence and brightfield images of dual reporter expression 48 hours following uniform seeding onto perpendicular GFP and mCherry patterns. Scale bars, 500μm. (G) Dual-positive miRFP and BFP masks, created using ImageJ image calculator AND function for BFP and miRFP signal above a threshold. Mask was superimposed onto fluorescence images and highlighted with a yellow border. Scale bars, 500μm.

**Figure S6.** (A) Fluorescence images of anti-GFP synNotch MyoD fibroblasts stained for α-actinin 7 days following seeding onto flat micromolded gelatin substrates in the presence of transglutaminase-conjugated GFP (100μg/mL). Scale bars, 500μm. Myotube alignment of α-actinin stained myotubes on GFP-conjugated isotropic (no topography) gelatin substrate, quantified by image analysis. Line plot represents angles of orientation distribution of 5 individual images from an individual sample. (B) Binary Mask and fluorescence images of anti-GFP synNotch MyoD fibroblasts seeded onto adhesion-restricted GFP patterned surfaces (500 and 200 µm rows) stained for α-actinin after 3 days. Scale bars, 500µm. Plot profile of normalized α-actinin staining intensity and GFP signal on adhesion-restricted 500µm and 200µm GFP row patterns. (C) Fluorescence images of nuclei staining for each pattern: 500µm curves, 200µm curves, 500µm rows, and 200µm rows. Samples were stained three days following uniform seeding onto GFP patterns. (D) Day 3 brightfield image of spatial myotube formation on 500µm width rows. Zoomed in region, indicated with a dotted white line, showing phenotype difference on-pattern (myotubes) vs off-pattern (fibroblasts). (E) Higher magnification fluorescence images of α-actinin stained cells on isotropic or 500µm row GFP patterns. Scale bars, 200µm. (F) Plot profiles of normalized α-actinin expression across 500 and 200µm rows on non-restricted GFP patterns. Green lines indicate the region containing GFP. (G) Staining validation of mouse α-actinin antibody on myotubes differentiated on GFP patterns, Day 3.

**Figure S7.** (A) Fluorescence images of anti-mCherry synNotch fibroblasts with inducible ETV2 – BFP expression in the presence of varying plate-dried mCherry concentrations (0-100µg/cm$^2$). BFP signal is shown in grayscale for visualization, images taken 1, 2, and 3 days following seeding. Scale bar, 200µm. (B) Normalized BFP intensity of anti-mCherry synNotch fibroblasts seeded on varying concentrations of plate-dried mCherry up to 3 days. Data represents mean ± s.d, n=3. (C) Staining validation of anti-mCherry synNotch fibroblasts seeded on 15µg/cm$^2$ mCherry with endothelial marker VEGFR2. Panel shows sample stained with or without an anti-VEGFR2 primary antibody. Inducible BFP reporter and nuclei stained with HSC NuclearMask Deep Red. Scale bars, 200µm. (D) Day 3 brightfield image of spatial endothelial-progenitor cell activation on vascular-like pattern. Dotted white line indicates the region of interest enhanced in the following panel. Scale bars, 1mm and 500µm, respectively. (E) Flow cytometry gating strategy to exclude debris (Gate 1), isolate singlets (Gate 2), and determine BFP-expressing cells (top) or VEGFR2 expressing cells (bottom).

**Figure S8.** (A) Fluorescence images of anti-GFP and anti-mCherry dual-lineage synNotch fibroblasts with orthogonally inducible MyoD-miRFP and ETV2 – BFP, respectively, expression in the presence of varying plate-dried mCherry concentrations (0-100µg/cm$^2$). BFP signal is shown in grayscale for visualization, images taken 1, 2, and 3 days following seeding. Scale bars, 200µm. (B) Normalized BFP intensity of dual-lineage synNotch fibroblasts seeded on varying concentrations of plate-dried mCherry up to 3 days. Data represents mean ± s.d, n=3. (C) Fluorescence images of anti-GFP and anti-mCherry dual-lineage synNotch fibroblasts with orthogonally inducible MyoD-miRFP and ETV2 – BFP expression in the presence of varying plate-dried GFP concentrations (0-100µg/cm$^2$). miRFP signal is shown in grayscale for visualization, images taken 1, 2, and 3 days following seeding. Samples from each GFP concentration were fixed and stained for α-Actinin following three days of culture. Scale bars, 200µm.

**Figure S9.** (A) Schematic of dual-fate embryonic fibroblasts expressing anti-GFP synNotch that activates MyoD and miRFP as well as an anti-mCherry synNotch that orthogonally activates ETV2 and BFP transgenes (anti-GFP→myoD+mCherry & anti-mCherry→ETV2+BFP) seeded onto GFP and mCherry droplet-patterned substrate. (B) Fluorescence images of GFP and mCherry droplet patterns (left), subsequent spatial reporter activation (center) taken 48 hours after uniform seeding of engineered dual-fate fibroblasts. Scale bars, 2mm. Green (within GFP pattern) and red (within mCherry pattern) borders indicate regions of interest with corresponding higher magnification brightfield and fluorescence images (right). Scale bars, 200µm. (C) Brightfield image of dual-lineage cells seeded on droplet pattern taken 48 hours after uniform seeding of engineered fibroblasts. Scale bars, 2mm. (D) Merged fluorescence images to demonstrate the effects of subtracting the GFP pattern (high GFP intensity excluded) in visualizing the α-actinin and VEGFR2 staining. Scale bars, 1mm. (E) Flow cytometry panel (X-Axis: VEGFR2-APC, Y-Axis: BFP) of dual-lineage cells seeded on 15μg/cm$^2$ plate-dried mCherry unstained (left) or stained for VEGFR2 (center) and dual-lineage cells seeded on 15μg/cm$^2$ plate-dried of both GFP and mCherry stained for VEGFR2 (right).
